# Comparison of Resources and Methods to infer Cell-Cell Communication from Single-cell RNA Data

**DOI:** 10.1101/2021.05.21.445160

**Authors:** Daniel Dimitrov, Dénes Türei, Charlotte Boys, James S. Nagai, Ricardo O. Ramirez Flores, Hyojin Kim, Bence Szalai, Ivan G. Costa, Aurélien Dugourd, Alberto Valdeolivas, Julio Saez-Rodriguez

## Abstract

The growing availability of single-cell data has sparked an increased interest in the inference of cell-cell communication from this data. Many tools have been developed for this purpose. Each of them consists of a resource of intercellular interactions prior knowledge and a method to predict potential cell-cell communication events. Yet the impact of the choice of resource and method on the resulting predictions is largely unknown. To shed light on this, we created a framework, available at https://github.com/saezlab/ligrec_decoupler, to facilitate a comparative assessment of methods for inferring cell-cell communication from single cell transcriptomics data and then compared 15 resources and 6 methods. We found few unique interactions and a varying degree of overlap among the resources, and observed uneven coverage in terms of pathways and biological categories. We analysed a colorectal cancer single cell RNA-Seq dataset using all possible combinations of methods and resources. We found major differences among the highest ranked intercellular interactions inferred by each method even when using the same resources. The varying predictions lead to fundamentally different biological interpretations, highlighting the need to benchmark resources and methods.

**Findings:**

- Built a framework to systematically combine 15 resources and 6 methods to estimate cell-cell communication from single-cell RNA data
- Cell-cell communication resources are often built from the same original databases and very few interactions are unique to a single resource. Yet overlap varies among resources and certain biological terms are unevenly represented
- Different methods and resources provided notably different results
- The observed disagreement among the methods could have a considerable impact on the interpretation of results

## 1. Introduction

The growing availability of single-cell RNA sequencing (scRNA-Seq) data is helping us improve our understanding of the cellular heterogeneity of tissues. Furthermore, Spatial Transcriptomics has recently emerged as a technology to measure gene expression while preserving the spatial distribution of cells in a sample, thus providing an unprecedented opportunity to decipher tissue architecture and organization ^1^. These advancements have in turn led to an increased interest in the development of tools for cell-cell communication (CCC) inference. CCC commonly refers to interactions between secreted ligands and plasma membrane receptors. This picture can be broadened to include secreted enzymes, extracellular matrix proteins, transporters, and interactions that require the physical contact between cells, such as cell-cell adhesion proteins and gap junctions ^2^. For simplicity, we refer to all of these events involving protein-protein interactions as CCC. CCC events are essential for homeostasis, development, and disease, and their estimation is becoming a routine approach in scRNA-seq data analysis ^3^.

A number of computational tools and resources have emerged that can be further classified as those that predict CCC interactions alone ^4–13^, and those that additionally estimate intracellular pathway activities related to CCC ^14–18^. Here, we focus on the former (Table 1). These CCC tools typically use gene expression information obtained by scRNA-Seq. In general, single cells are clustered by their gene expression profile and cell type identities are assigned to the clusters based on known gene markers. Then, CCC tools can predict intercellular crosstalk between any pair of clusters, one cluster being the source and the other the target of a CCC event. CCC events are thus typically represented as a one-to-one interaction between a ‘transmitter’ and ‘receiver’ protein, expressed by the source and target cell clusters, respectively. The information about which transmitter binds to which receiver is extracted from diverse sources of prior knowledge. Roughly, CCC tools then estimate the likelihood of crosstalk based on the expression level of the transmitter and the receiver in the source and target clusters, respectively. Every tool has two major components: a resource of prior knowledge on CCC (interactions), and a method to estimate CCC from the known interactions and the dataset at hand. Most tools have been published as the combination of one resource and one method, but in principle any resource could be combined with any method.

**Table 1.** The tools included in the framework. Each tool uses a resource and a method with a specific scoring system. Each method considers expression at the cell cluster level, and all of the scoring systems presented here are based on the expression of transmitters and receiver genes in the source and target cells, respectively. CellPhoneDBv2 algorithm was included via its implementation in Squidpy ^24^. For further details, check the original references.

| Tool | Resource | Methods' scoring systems |
| --- | --- | --- |
| CellChat<sup>13</sup> | CellChatDB | 1) **Probability** - based on the expression of differentially expressed transmitter and receiver genes and their mediators, calculated with the law of mass action<br>2) **P-values†** - significance identified via permutation of cell cluster labels and recalculating the probabilities for each cell pair and each transmitter-receiver interaction |
| Squidpy#<sup>24</sup><br>(CellPhoneDB<sup>7,25,26</sup>) | OmniPath or CellPhoneDB | 1) **Truncated Mean** - average expression of transmitter and receivers, the minimum expression (by default) of heteromeric complex of subunits<br>2) **P-values†** - significance identified via permutation of cell cluster labels to determine a null distribution of means for each receiver-transmitter interaction |
| Connectome<sup>9</sup> | Ramilowski | 1) **weight_norm** - derived via the product (by default) of the normalized expression of transmitter and receiver genes<br>2) **weight_scale†** - derived from the function (mean, by default) of the z-scores of the transmitter and the receiver, scaled according to cell cluster specificity |
| iTALK<sup>5</sup> | iTALK | 1) **Expression Z-score products** - based on the differentially expressed transmitter and receiver genes between clusters |
| NATMI<sup>10</sup> | ConnectomeDB 2020 | 1) **Mean-expression edge weight** - transmitter and receiver gene expression product<br>2) **Specificity-based edge weight†** - the mean expression of the transmitter and receiver are divided by the sum of the means of the same transmitters/receivers across all cell clusters |
| SingleCellSignalR<sup>11</sup><br>(SCA) | LRdb | 1) **LRScore** - a regularized score calculated using the squared expression of the transmitter and receiver (sqTRE) divided by sum of the mean of the count matrix and sqTRE |
| † Explicitly measures cell-cluster specific communication (referred as “cluster-specific measures”)<br># Here we refer to the re-implementation of the CellPhoneDB method as ‘Squidpy’, even though Squidpy is a spatial transcriptomics framework with a much broader range of functionalities. |  |  |

Despite the aforementioned common premises to explore CCC events, each tool uses a different method, such as permutation of cluster labels, regularizations, and scaling, to prioritize interactions according to the input datasets (Table 1). In turn, these different approaches result in diverse scoring systems that are difficult to compare and evaluate. The difficulties are further exacerbated by the lack of an appropriate gold standard to benchmark the performance of CCC methods ^3, 19^. Nevertheless, different strategies have been used to indirectly evaluate the methods’ performance, including a presumed correlation between CCC activity and spatial adjacency ^13, 18^, recovering the effect of receptor gene knockouts ^18^, robustness to subsampling ^13^, agreement with proteomics ^11^, simulated scRNA-Seq data ^8^, and the agreement among methods ^9, 11, 13, 18^.

The available prior knowledge resources, largely composed of ligand-receptor, extracellular matrix, and adhesion interactions, are typically distinct but often show partial overlap ^2, 20^. Some of these resources also provide additional details for the interactions such as information about protein complexes ^2, 7, 13, 21, 22^, subcellular localisation ^2, 13^, and classification into signalling pathways and categories ^13, 21^ (Supp. Table 1). CCC resources are often manually curated and/or built from other resources, with varying proportions of expert curation and literature support ^2, 20^. Some databases gather and harmonize the information contained in the individual resources ^2^. Despite the fact that CCC inference is constrained by the prior knowledge used, the impact of resource choice is largely unexplored, with the only exception, to our knowledge, of a descriptive comparison of 4 resources with one method ^20^. It remains thus unclear how the choice of resource and method affects the results and thereby the biological interpretation of the scRNA-seq data.

Here, we systematically compared all combinations of 15 resources and 6 CCC methods (Figure 1). First, we explored the degree of overlap among resources and whether certain resources are biased toward specific biological terms, such as pathways and functional cancer states. Then, we analysed how different combinations of resources and methods influence CCC inference, by decoupling the methods from their corresponding resources. In particular, we explored their impact on the predicted CCC using a publicly available colorectal cancer scRNA-Seq dataset ^23^. Our framework, available at https://github.com/saezlab/ligrec_decoupler, establishes a uniform interface to all the resources and methods in any combination. We see this work as a platform for further analyses, benchmarks, and method development, and we invite all interested parties to join us in this endeavour.

**Figure 1.**
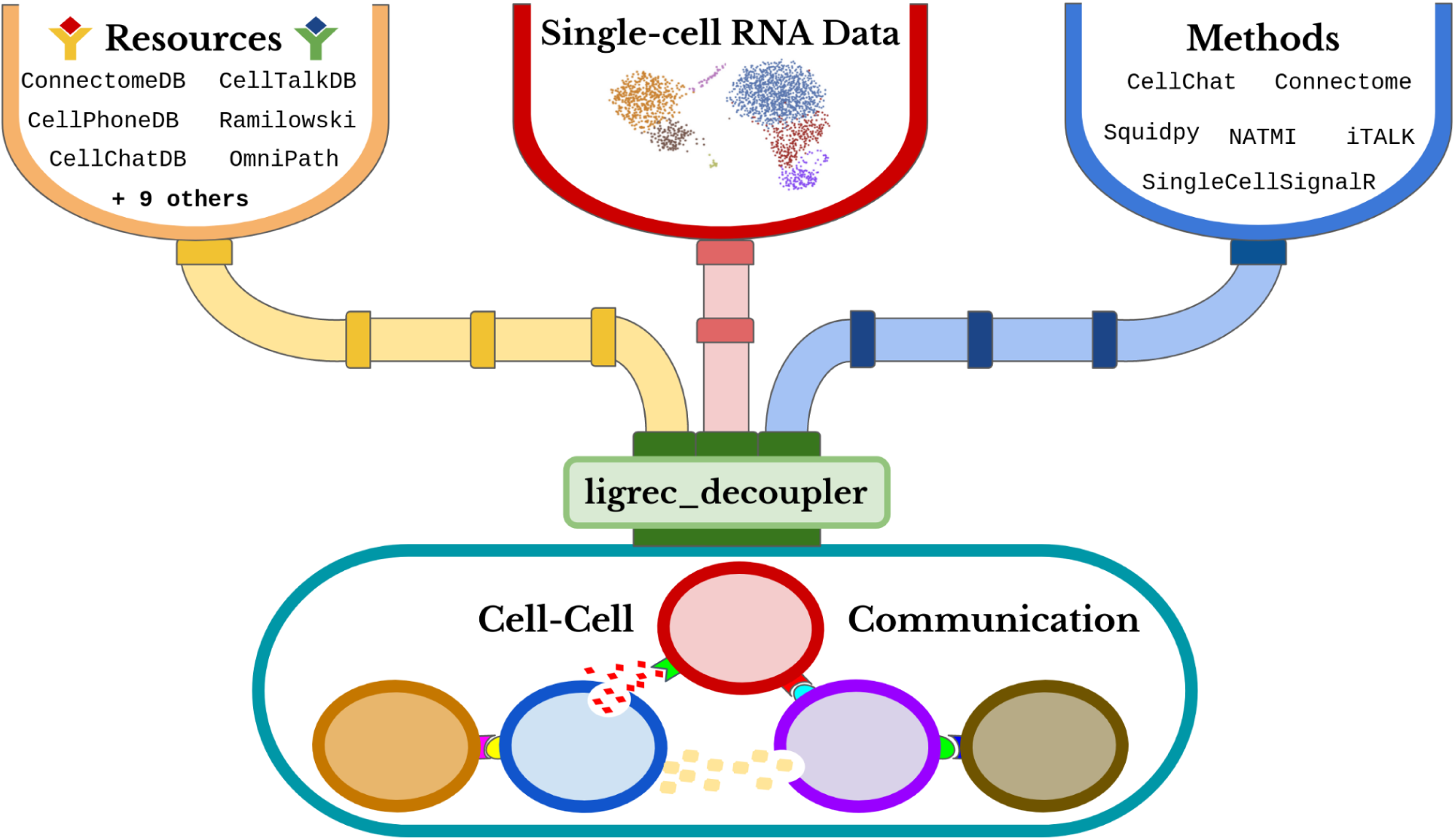
The Cell-Cell Communication Framework.

## 2. Results

### 2.1 Resource Uniqueness and Overlap

To investigate the lineages of CCC resources, we manually gathered information about the origins of each resource. Many of these resources share the same original data sources, including general biological databases such as KEGG ^27, 28^, Reactome ^29^, and STRING ^30^ (Figure 2A). Moreover, interactions from Guide to Pharmacology ^31^, CellPhoneDB ^7^, and in particular Ramilowski ^32^, were incorporated into subsequently published resources. All these resources are integrated into OmniPath’s CCC resource ^2^, along with additional CCC interactions from other sources (e.g. SIGNOR ^33^, Adhesome ^34^, SignaLink ^35^, and others). We filtered the OmniPath CCC interactions by quality (4.1 Methods).

**Figure 2.**
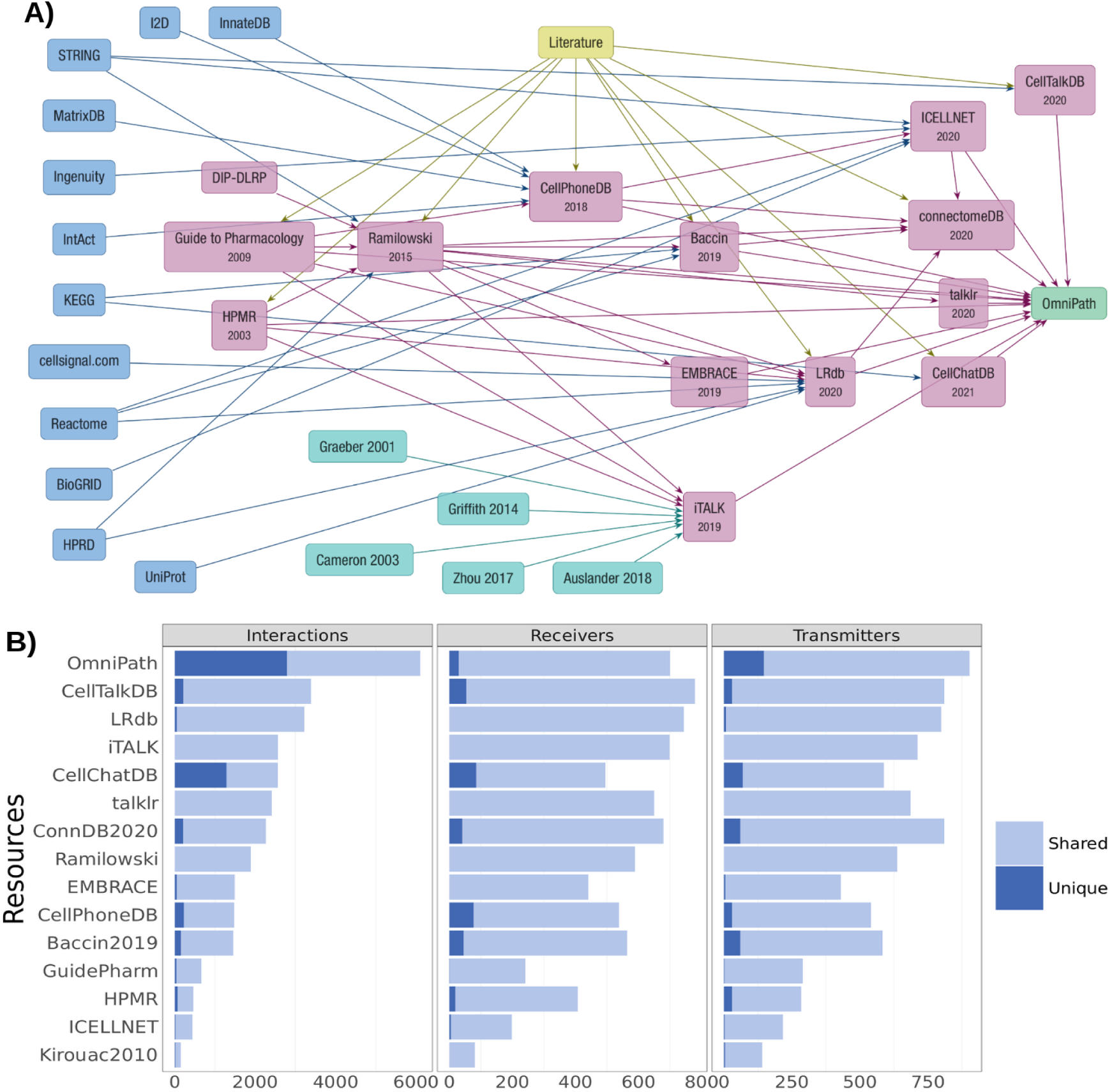
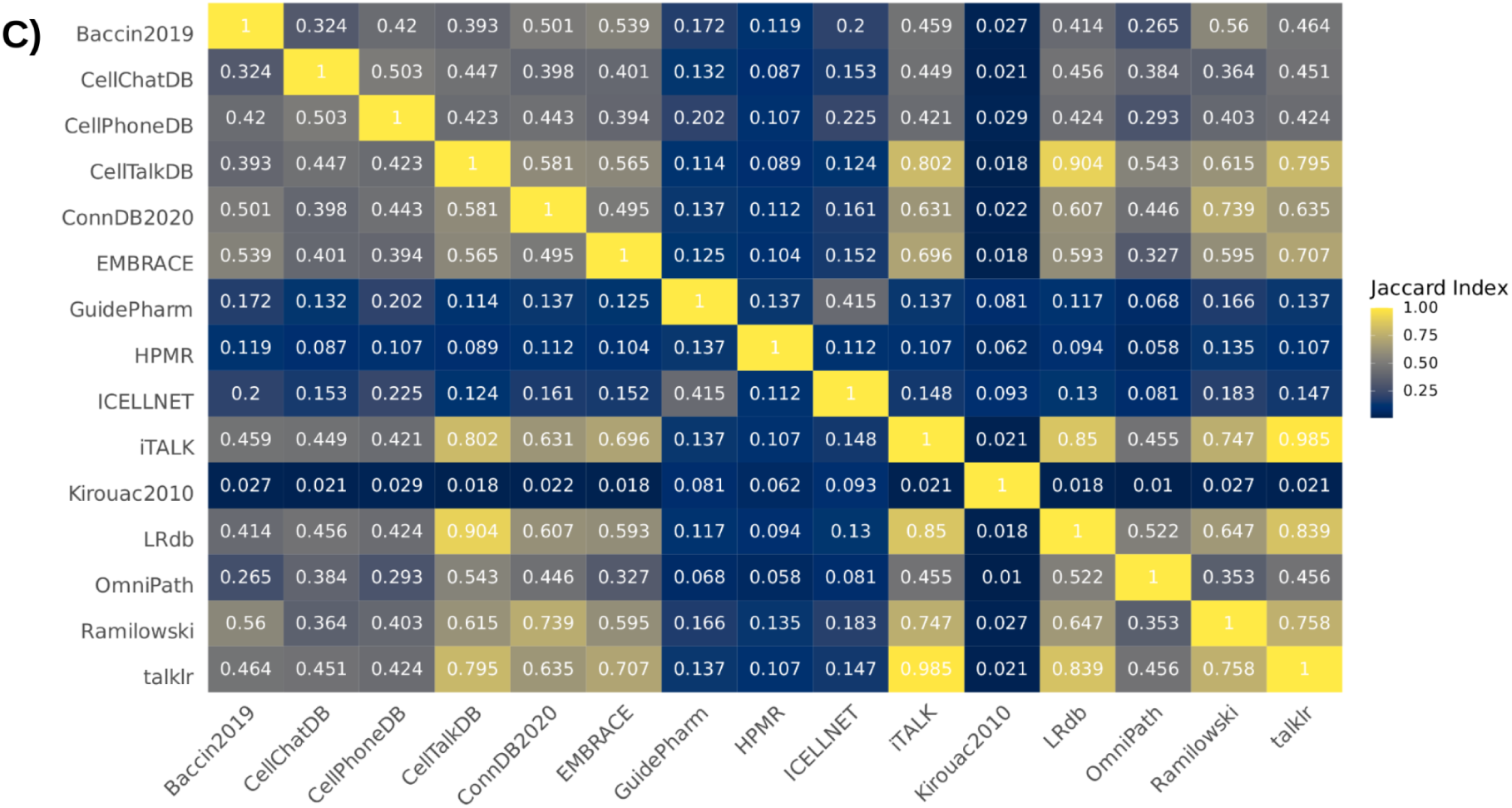
Dependencies and overlap between CCC resources. A) The lineages of CCC interaction database knowledge. General biological knowledge databases, CCC-dedicated resources used in this work, Literature curation, resources included in iTALK, and OmniPath are in blue, magenta, yellow, cyan, and green respectively. Arrows show the data transfers between resources. B) Shared and Unique Interactions, Receivers and Transmitters. C) Similarity between the interactions from different resources (Jaccard Index).

As a consequence of their common origins, we noted limited uniqueness across the resources, with mean percentages of 4.6 unique receivers, 5.3 unique transmitters, and 16.8% unique interactions, for all resources (Figure 2B; Supp. Table 1). OmniPath and CellChatDB ^13^ had the largest degree of uniqueness, with 4, 16, and 46% for OmniPath and 17, 12, and 50% for CellChatDB in terms of receivers, transmitters, and interactions, respectively. Despite the sparse uniqueness among the resources, the pairwise overlap between them varied (Figure 2C; Supp. Figure 1). Particularly high similarity was observed between CellTalkDB ^20^, ConnectomeDB ^10^, talklr ^36^, iTALK ^5^, LRdb ^11^, and Ramilowski (Figure 2C; Supp. Figure 2). The aforementioned resources, together with OmniPath, contained on average more than 65% the interactions present in the other resources (Supp. Figure 3), largely explained by each including a large proportion (>80%) of the interactions present in Ramilowski. CellChatDB, CellPhoneDB and Baccin ^22^ showed limited similarity with other resources, as each included ∼45% of the interactions present in any other resource, on average. These latter resources include protein complexes, which were dissociated and treated as distinct protein subunits in our resource analyses. The smaller resources ICELLNET ^12^, Guide to Pharmacology, HMPR ^37^ and Kirouac2010 ^38^ were most dissimilar with the remainder of the resources and included on average only 21, 28, 17, and 7% of the interactions present in the other resources, respectively. The similarity among the resources was generally higher when considering transmitters, and receivers in particular (Supp. Figure 1-3).

In summary, our results indicate that many of the transmitters, receivers, and interactions are not unique to any single resource, due to their common origins. However, different resources include varying proportions of the collective CCC prior knowledge.

### 2.2 Resource Prior Knowledge Bias

Since CCC inference methods rely on prior knowledge to estimate intercellular communication events, the choice of resource and any potential bias in it is expected to impact the results. We therefore explored whether the coverage of interactions in the resources is biased toward specific subcellular locations or functional categories when compared to the collection of all resources.

#### 2.2.1 Subcellular Localisation

We obtained protein subcellular localisation annotations from OmniPath ^2^, which combines this information from 20 resources. We then matched the localisations to receivers and transmitters from each resource with the aim to assess the localisation profile of different resources. On average 90% of transmitters and 79% of receivers were annotated as secreted and transmembrane proteins, respectively (Supp. Figure 4). We further used the localisations of transmitters and receivers to categorize the interactions as secreted or direct-contact signaling. We reasoned that, interactions between transmitters annotated as secreted and receivers annotated as membrane-bound represent solute mediated (secreted) signalling events. On the contrary, an interaction between two membrane-bound proteins requires direct contact between cells. Building on this, we observed that all resources were predominantly (74% on average) composed of interactions associated with secreted signalling, while direct-contact signalling constituted a substantially smaller (16% on average) proportion of interactions (Figure 3A; Supp. Figure 5). Interactions categorized as neither secreted nor direct-contact were labeled as ‘Other’ and made up the remainder of the interactions (Supp. Note 1). The proportions of secreted and direct-contact signalling varied between resources, as some of them, such as Baccin, ConnectomeDB, CellPhoneDB, HPMR, and OmniPath had an over-representation of direct-contact signalling when compared to the collective, while the opposite was noted for the case of secreted signalling (Figure 3B). Direct contact interactions were particularly under-represented in Guide to Pharmacology (4%), which was more focused on secreted signalling (87%). CellChatDB showed an overrepresentation of interactions matched to the category Other.

**Figure 3.**
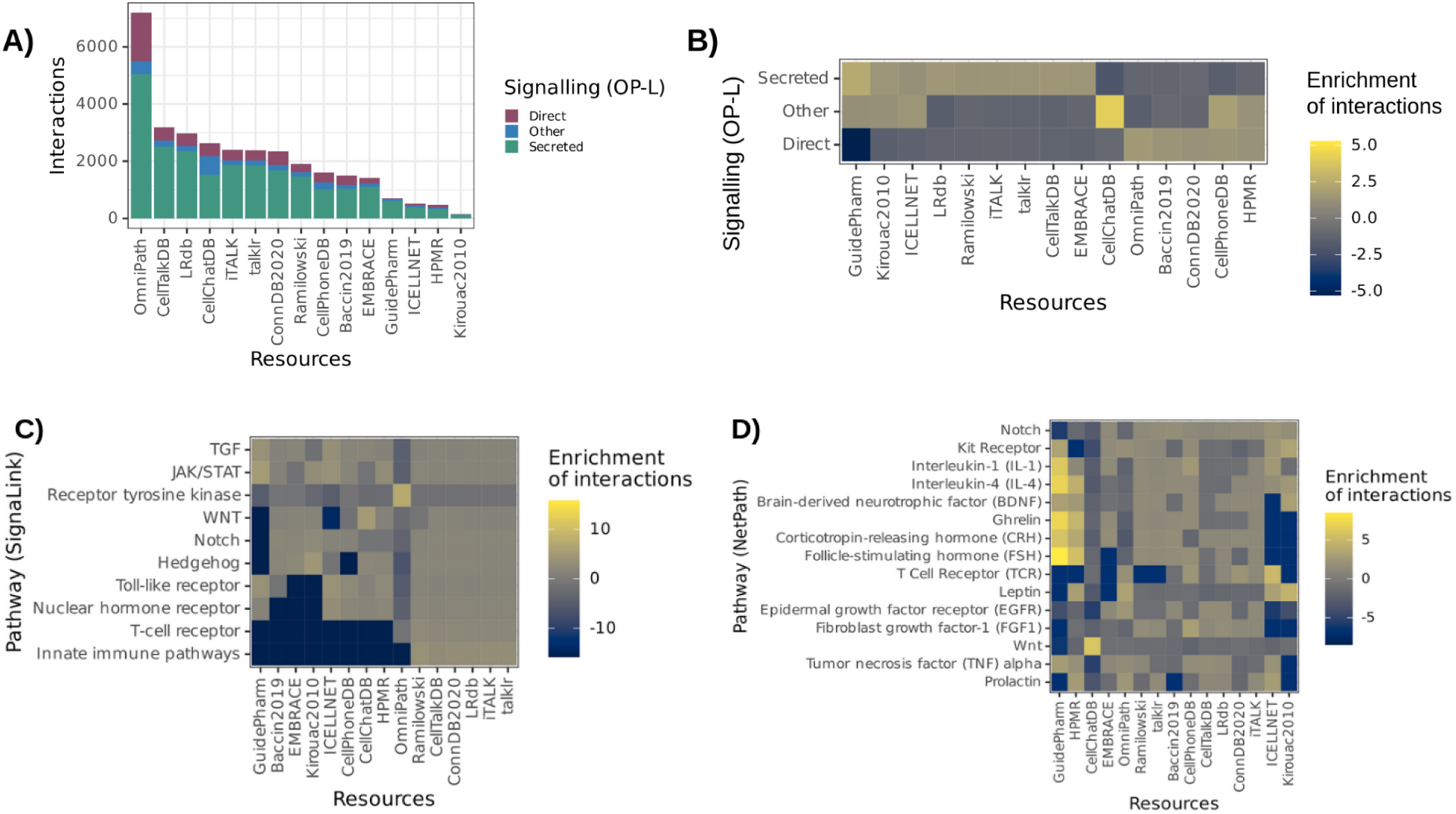
Localisation and Functional term abundance in CCC resources. A) Number and B) Relative abundance of signalling categories based on OmniPath-derived protein locations (OP-L) by resource. Relative abundance of C) SignaLink and D) NetPath annotations matched to interactions from each resource.

Our results suggest that localisations of transmitters and receivers were largely uniformly distributed and that secreted signalling was predominant across all resources. Yet, differences were noted between the relative abundance of secreted and direct-contact signalling interactions.

#### 2.2.2 Functional Term Enrichment

To examine whether specific pathways and biological functions are unevenly represented in specific resources, we matched the interactions, receivers and transmitters from each resource to well-known pathways and functional categories from SignaLink ^35^, NetPath ^39^, CancerSEA ^40^, and HGNC ^41^.

We observed that the Receptor tyrosine kinase (RTK), JAK/STAT, TGF, and WNT pathways covered the largest proportions of interactions matched to SignaLink, with analogous results observed for receivers and transmitters (Supp. Figure 6). The interactions from Ramilowski, ConnectomeDB, LRdb, iTALK and talklr showed a similar pattern, which can be explained by the high overlap of these resources. On the contrary, interactions associated with innate immune pathways and T-cell receptor categories were under-represented in Guide to Pharmacology, Baccin2019, EMBRACE, Kirouac2010, ICELLNET, CellPhoneDB, and HMPR (Figure 3C). The innate immune pathway category was also diminished in OmniPath. In contrast, when we used NetPath instead of SignaLink to define the T-cell receptor pathway, the under-representation in Baccin2019 and OmniPath was not observed, and an over-representation was instead noted for ICELLNET and CellPhoneDB (Figure 3D; Supp. Figure 7). Moreover, we observed a considerable over-representation for the RTK pathway in OmniPath. The Signalink WNT pathway was under-represented in ICELLNET and Guide to Pharmacology, while for the NetPath WNT pathway this was only true for Guide to Pharmacology. In contrast, CellChat showed a relative abundance for both the SignaLink and NetPath WNT pathways. These observations for the WNT pathway were further supported by the relative abundance of HGNC (Supp. Figure 8). Functional cancer cell states from CancerSEA were also unevenly represented in sets of receivers and transmitters across the resources (Supp. Figure 9). For example, Cell Cycle, DNA repair, and DNA damage states were over-represented in LRdb. Hence, our results indicated heterogenous biases towards certain pathways and categories across the different CCC resources.

### 2.3 Agreement in CCC predictions in a Colorectal Cancer data set

To estimate the relative agreement between CCC methods and the importance of resources, we developed a framework to decouple tools from their inbuilt resources. We chose a well-annotated colorectal cancer (CRC) scRNA-Seq dataset ^23^ with 65,362 cells from a heterogeneous cohort of 23 Korean CRC patients. The 38 cell annotations in the dataset included stromal, immune, tumour and healthy epithelial cell types/states, as well as 3 unknown subtypes of Myeloid, B and T cells. We focused on the interactions between tumour cells subclassified by their resemblance of CRC consensus molecular subtypes (CMS) and immune cells from tumour samples (Supp. Table 3), reasoning that this subset of cell types represents a complex example where CCC events are known to have an important role. In addition to the 15 CCC resources reported in the descriptive resource analysis (Supp. Table 2), we also included the default or inbuilt resource for each of the tools, except Squidpy (Table 1), as well as a reshuffled control resource (4.3 Framework).

#### 2.3.1 Interaction overlap

We then used each method-resource combination to infer CCC interactions, assuming that different methods should generally agree on the most relevant CCC events for the same resource and expression data. To measure the agreement between method-resource combinations, we looked at the overlap between the 500 highest ranked interactions as predicted by each method. Whenever available, author recommendations were used to filter out the false-positive interactions (4.4 Method-Resource Specifics). Our analysis showed considerable differences in the interactions predicted by each of the methods regardless of the resource used, as the mean Jaccard index per resource ranged from 0.01 to 0.06 (mean = 0.024) when using different methods. These large discrepancies in the results were further supported by the pairwise comparisons between methods using the same resource, with mean Jaccard indices ranging from 0.063 (CellChat-SingleCellSignalR) to 0.110 (Connectome-NATMI). The overlap among the top predicted interactions was slightly higher when using the same method but with different resources, as Jaccard indices ranged from 0.113 to 0.203 per method (mean = 0.167) (Supp. Figure 11). Consequently, the highest ranked interactions for each method-resource combination largely showed stronger clustering by method than resource (Figure 4), with similar results observed when considering the highest ranked 100, 250, and 1000 interactions (Supp. Figure 10). In particular, method-resource combinations involving Squidpy, SingleCellSignalR, and Connectome clustered exclusively by method, suggesting that the overlap between these combinations occurs predominantly when using the same method regardless of the resource (Supp. Figure 11). The combinations involving NATMI also clustered by method, with the only exceptions being the Kirouac2010 ^38^ and ICELLNET ^21^ resources, which were the smallest resources (Supp. Table 2).

**Figure 4.**
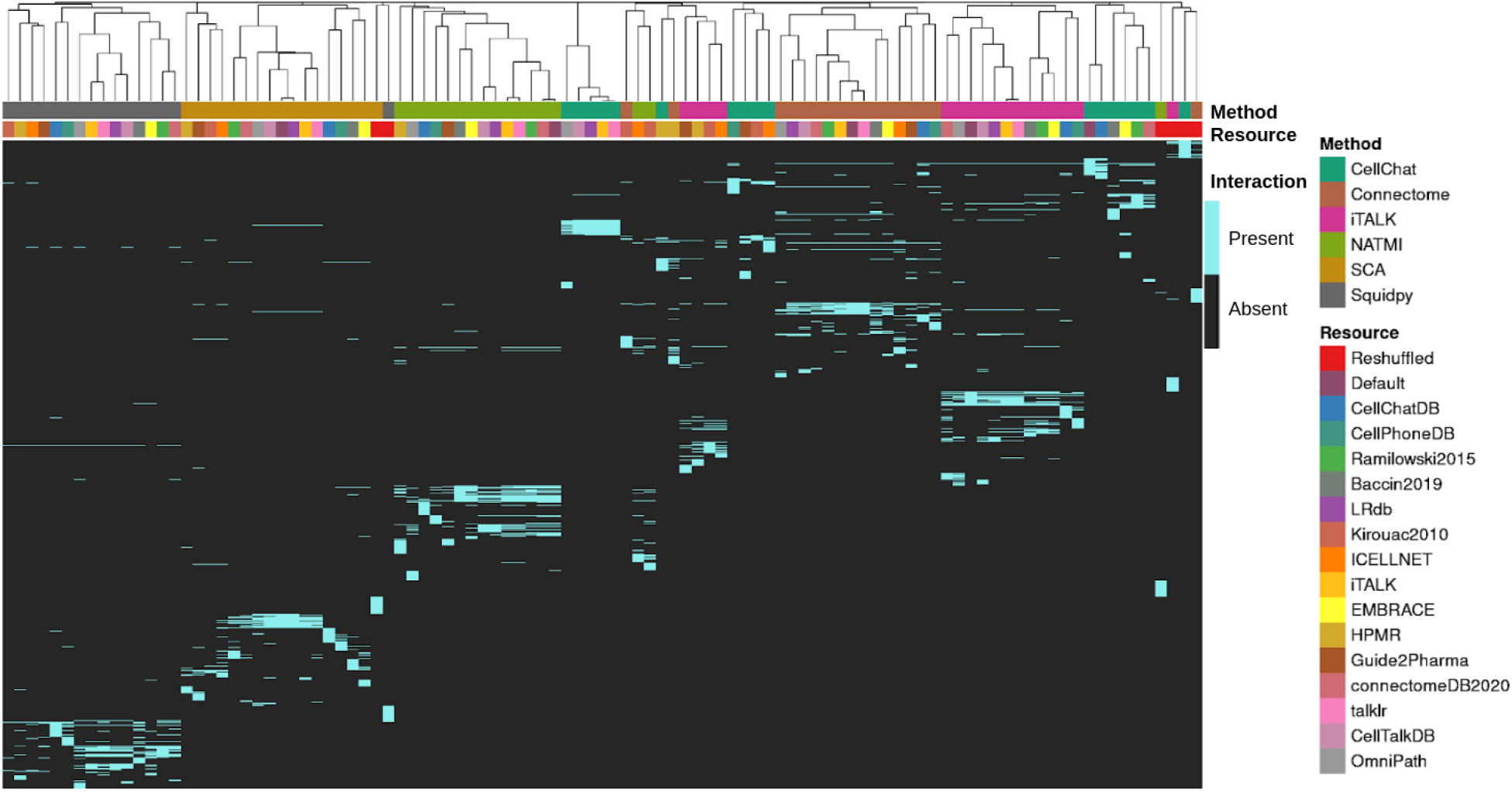
Overlap in the 500 highest ranked CCC interactions between different combinations of methods and resources. Method-resource combinations were clustered according to binary (Jaccard index) distances. SCA refers to the SingleCellSignalR method.

Moreover, CellChat and the CellPhoneDB (Squidpy) methods ^7, 24^, account for heteromeric transmitter-receiver complexes, as such we examined the proportion of complex-containing interactions for these methods using the complex-containing resources (Supp. Note 2). This analysis showed that the proportion of complexes among the highest ranked hits was 2-23% for CellChat and 10-38% for Squidpy, largely reflecting the relative complex content in each resource.

Our results suggest that the overlap between methods when using the same resource was low (Supp. Figure 12). This was largely supported by the analysis of two additional scRNA-Seq data of cord blood mononucleated cells and pancreatic islet cells (Supp. Figure 13), even though we observed slightly higher agreement between methods in the former dataset. The overlap when using the same method with different resources, albeit higher than that between different methods, was also modest (Supp. Figure 14). Hence, our results indicate that both the method and the resource had a considerable impact on the predicted interactions.

#### 2.3.2 Communicating cell types

Next, we asked whether the discrepancies observed between the methods stem from the differences in the cell types inferred as most active in terms of CCC interactions. To this end, we used the 500 highest ranked interactions to examine the cell type activities, defined as the proportion of interactions per cell type, separately as a source and a target of CCC events (Figure 5). The results largely reiterated our observations from the CCC interaction overlap analysis above, as each method largely clustered by itself, regardless of the resource used, including the reshuffled resource. These results were further supported by the average interaction ranks per communicating pairs of clusters, as again the method-resource combinations largely grouped by method (Supp. Figure 15). We reasoned that the observed disagreement in regards to the most actively communicating cell types was likely caused by the methods’ distinct approaches to handle cell cluster specificity. We thus performed a complementary analysis using the alternative, non cell-type specific scoring systems of the methods. The higher agreement in this case suggested that these different approaches are indeed in part responsible for the disagreement (Supp. Note 3).

**Figure 5.**
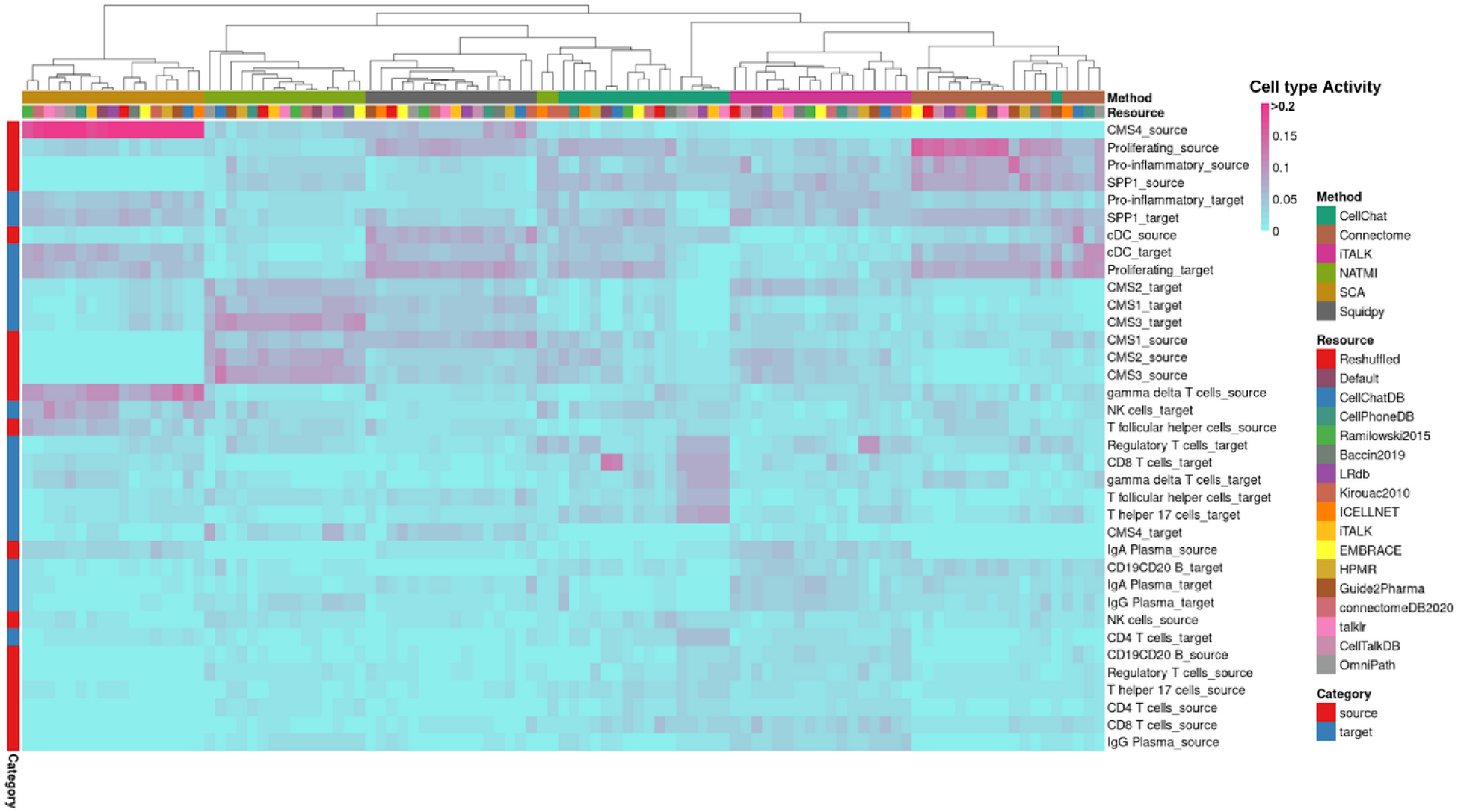
Activity per Cell type, inferred as the proportion of interaction edges that stem from Source Cell clusters or lead to Target Cell clusters in the highest ranked interactions.

We further argued that the choice of method can have a major impact on the interpretation of CCC results. For example, regardless of the resource, SingleCellSignalR predicted CMS4-like cells to be a major source of signalling within the system, which was in disagreement with the majority of method-resource combinations (Figure 5). Nevertheless, given that CMS4 is characterized by the exertion of immunosuppressive pressure on immune cells via stromal cells ^42^, it can be argued that SingleCellSignalR appropriately recognized CMS4 as the most active source of signalling. Regardless of the resource, NATMI highlighted CMS1-, CMS2-, and CMS3-like tumour cells as both major sources and targets of signalling, while the inferred activity of immune cells was overall sparse. NATMI’s predictions were supported by CMS2 and CMS3 tumor subtypes being associated with having low immune and inflammatory molecular signatures ^43^. Yet, this is not expected to be the case for CMS1-like cells, as CMS1 tumors are characterized by the infiltration of cytotoxic lymphocytes ^42, 43^. Moreover, CD4+ T cells and CD8+ T cells were the two most abundant immune cell types within the dataset (Supp. Table 3), but they were estimated to be among the least active cell types in the system across most method-resource combinations. Secreted phosphoprotein 1 (*SPP1*)^+^ macrophages were observed to be important sources and targets of cell-cell communication in the system across most method-resource combinations. This observation is largely supported by the enrichment of SPP1+ macrophages in tumour tissues, and their potential key role in tumor progression and immune suppression ^23^.

The analysis of activities per cell type largely reiterated the results from the interaction overlap analysis, particularly as each method largely clustered by itself, regardless of the resource. As a consequence, the disagreement between the methods in which cell types are the most active is expected to have a major impact on the biological interpretation of CCC communication predictions.

## 3. Discussion

The growing interest in CCC inference has led to the recent emergence of multiple methods and prior knowledge resources dedicated to studying intercellular crosstalk. To shed light on the impact of the choice of method and resource on the inference of CCC events, we built a framework to systematically combine 15 resources and 6 methods. We used this framework to describe in detail the content of the different resources and to estimate cell-cell communication from scRNA-Seq in a colon cancer case study. Our results suggest that both the method and resource can considerably impact CCC inference.

### 3.1. Resource Overlap and Bias

Despite their common origins, different resources cover varying proportions of the collective prior knowledge. Particularly, a large share of the observed overlap among resources stemmed from the inclusion of Ramilowski ^32^ into other resources. Moreover, across the resources, the WNT, RTK, T-cell receptor and Innate immune pathways, among others, were present in varying proportions. The high abundance of interactions associated with the RTK pathway in OmniPath could be due to the ∼1,600 expert curated RTK ligand-receptor interactions from SIGNOR ^33^ and the large size of RTK pathway in SignaLink ^35^. The results presented here highlight an inherent limitation of knowledge-based inference, and hence of CCC methods, as any prior knowledge resource has its own biases and only represents a limited proportion of biological actuality. Consequently, these inherent limitations should be kept in mind for the interpretation of CCC predictions.

### 3.2. Impact of Methods and Resources

As a further step, we carried out a systematic analysis of the impact of resources and methods on CCC inference results using a public colorectal cancer dataset ^23^. Although possibly over-simplistic, our binary overlap assessment enabled the direct comparison of the diverse scoring systems of the methods. We found that both resources and methods had a considerable effect on the predicted interactions, and the impact of methods outweighed that of the resource.

A potential explanation for the disagreement among the methods could be the distinct approaches they use to identify the most relevant interactions (Table 1). A common assumption among the methods is that cluster-specific interactions are more informative than those related to multiple clusters ^7, 9, 10, 13^. An experimental proof of this assumption and an evaluation of the distinct approaches is yet to be carried out. By focusing on the cluster-specific interactions in the dataset, these methods report the most specifically-interacting cell types ^11^, rather than the most actively communicating ones. Hence, the predicted CCC events typically do not capture processes that are common between multiple cell types. As an example, CD4+ T cells and CD8+ T cells, the two most abundant immune cell types found within tumours, were assigned a low CCC communication activity. In terms of the agreement in CCC event predictions we found that only a few biological patterns are robust across many methods, namely the SPP1+ macrophages have been predicted to be main players of CCC signalling, supported by their enrichment in CRC tumour tissues ^23^ and frequent association with pro-metastatic role ^44, 45^. Nevertheless, we acknowledge that the pooled analysis of the CRC dataset largely limits the interpretation of results and warrants the future analysis of the same dataset on a per-patient/phenotype basis. Collectively, these results suggest that the common practice to highlight the most actively communicating cell clusters based on the CCC inference ^23, 46^ should be considered with caution.

### 3.3. Previous Comparisons and limitations of our integration

Interestingly, our results did not recover some of the previously reported agreement between tools ^9–11, 13^. Contrary to previous comparisons^11^, we saw little overlap between the SingleCellSignalR, CellPhoneDB and iTALK methods. Furthermore, despite their relatively similar approaches, the limited agreement between CellChat and CellPhoneDB was not observed here ^13^. It is to note that we used CellChat’s probabilities instead of p-values to obtain the highest ranked interactions. These probabilities do not deliberately reflect cell cluster specificity in regards to the inferred interactions^13^. However, our results support the previously observed low agreement between CellChat and both iTALK and SingleCellSignalR^13^.

Some methods, namely CellChat ^13^ and the CellPhoneDB algorithm ^7^, as well as resources, such as Baccin, CellChatDB, CellPhoneDB, and ICELLNET, take protein complexes into account. This largely complicates the conversion of the resources and hence the comparison with methods and resources which do not consider complexes. Furthermore, CellChat, and hence CellChatDB, goes a step further than other methods and resources, as it considers interaction mediator molecules, which are absent in the remainder of the resources ^13^. Thus, even though any resource can be used with any method, we acknowledge that some combinations put certain methods at a disadvantage.

### 3.4. CCC Inference Assumptions and Benchmarking

Our results further point to certain limitations of the CCC inference methods. In particular, CCC events are mainly predicted based on the average gene expression at the cluster or cell type/state level. Such an assumption inherently suggests that gene expression is informative of the activity of transmitters and receivers. However, gene expression provided by scRNA-Seq is typically limited to protein coding genes and the cells within the dataset, and hence does not capture secreted signalling events driven by non-protein molecules or long-distance endocrine signalling events. Further, CCC inference from scRNA-Seq data assumes that the product of the gene expression of a transmitter and a receiver is a good proxy for their joint activity, and thus does not consider any of the processes preceding transmitter-receiver interactions, including protein translation and processing, secretion, and diffusion.

We therefore believe that it is essential to establish a benchmark to comprehensively assess the predictive power of CCC methods. However, a gold standard for benchmarking is currently not available and the biological ground truth is largely unknown ^3, 19^. The field needs to identify experimental settings capable of establishing the biological ground truth. So far, intercellular interactions were mainly supported by the spatial colocalization of proteins and the functional deregulation of intracellular signalling ^47^, and the physical-interaction of cell types ^48^. Yet these approaches are only applicable for the post-hoc and indirect validation of CCC interactions. Thus, until an experimental gold standard becomes available, simulated datasets might be used instead. However, any in silico benchmark is by definition only a simplified approximation of reality, with its own biases ^49^. To our knowledge, appropriate benchmarks for resources and methods used in CCC inference are yet to be defined, although some proposals exist ^19^, that we elaborate on in Supp. Note 4.

### 3.5. Conclusion

Considerable efforts have been made to develop CCC inference, and we believe that further advancements will be key for the systems-level analysis of single-cell data. This will likely further increase by the rapidly emerging spatial transcriptomics ^1^ and single-cell proteomics ^50^, and the future applications of CCC inference approaches to interspecies communication ^51, 52^. Acknowledging the limitations of our work, we believe that it points at the interpretation inconsistencies that could arise as a consequence of the method and resource of choice. We thus regard the results and comparative framework presented here as steps towards an understanding of the strengths and weaknesses of CCC methods, and thereby towards their improvement. Future developments of this work will include extending the number of datasets in the comparative analysis as well as the benchmark of methods and resources. As such, we here extend an open invitation to all interested parties willing to join us in this endeavour.

## 4. Methods

### 4.1. Descriptive analysis of resources

The connections between resources shown in the dependency plot were manually gathered from the publications and the web pages of each CCC resource.

The CCC resources used in the analyses were queried from the OmniPath database^2^. The contents of the resources are identical to their original formats, apart from minor processing differences (Supp. Table 2).

The OmniPath CCC resource is a composite resource which contains interactions from all of the CCC dedicated resources compared here, along with some additional resources^2^. OmniPath’s interactions were filtered according to the following criteria: i) we only retained interactions with literature references, ii) we kept interactions only where the receiver protein was plasma membrane transmembrane or peripheral according to at least 30% of the localisation annotations, and iii) we only considered interactions between single proteins (interactions between complexes are also available in OmniPath). OmniPath’s intra- and intercellular components are both available via the OmnipathR package (https://github.com/saezlab/OmnipathR).

We defined unique and shared interactions, receivers and transmitters between the CCC resources if they could be found in only one or at least two of the resources, respectively. We used pheatmap ^53^ and UpSetR ^54^ to generate the heatmaps and upset plots, respectively.

To identify uneven distributions of transmitters, receivers, and interactions toward biological terms or protein localisations, we used Fisher’s exact test to compare each individual resource to the collection of all the resources. We obtained protein localisations from OmniPath which collects this information from 20 databases^2^. Then we kept consensus protein localisations where at least 50% of the annotations agreed. We classified CCC interactions using the localisation combinations of proteins involved in the interactions, which included secreted, plasma membrane peripheral and transmembrane proteins. Interactions, receivers and transmitters were independently matched to the 10 pathways from SignaLink ^35^, and the 15 largest categories from CancerSEA ^40^, HGNC ^41^, and NetPath ^39^. Each of the aforementioned general functional annotation databases was also obtained via OmniPath. In case of signalling pathway databases (SignaLink and NetPath), we focused on the enrichment of annotations matched to interactions, while for the functional state databases (CancerSEA and HGNC), we presented the merged sets of transmitters and receivers matched to functional categories. Annotation matches for transmitters and receivers were examined independently using the aforementioned functional annotation databases, but they were not the focus of discussion presented here. Our approach allowed the same protein or interaction to be matched to multiple pathways or functional categories from the same database.

To enable a comparison of annotations across resources, we expanded protein complexes from Baccin2019 ^22^, CellChatDB ^13^, CellPhoneDB ^7^, and ICELLNET ^10^.

### 4.2. Single-cell Transcriptomics data

The processed single cell RNA-Seq data ^23^ for 23 Korean colorectal cancer patients is available at GSE132465. The analysis presented here focused on the CCC interactions between colorectal cancer subtypes and immune cells, and the remainder of cell types, including unknown immune cell subtypes, were filtered out. This resulted in a subset of 18 cell types and 42,544 cells. We kept the original subtype labels, reformatted the names to work with each CCC method (Supp. Table 3), and sparsified the counts into a Seurat^55^ object.

The labelled scRNA-Seq data for pancreatic islet ^56^ and cord blood mononuclear cells ^57^ were obtained via SeuratData, normalized with Seurat ^55^, and used for CCC inference without any further formatting and filtering.

### 4.3. Framework

For the method-resource comparison, we used Seurat ^55, 56^ objects which were converted into the appropriate data format when calling each method. We used the recommended conversion method or wrapper whenever available.

The resources were obtained from OmniPath and then converted to the appropriate format for each method. A reshuffled version of ConnectomeDB2020 was generated with BiRewire ^58^ and referred to as the reshuffled control resource. Each tool was run with its default or inbuilt resource, except Squidpy. The Default resource of Squidpy’s **ligrec** function is OmniPath, which is already part of our benchmark set. The framework enabling the use of any resource and method combination, as well as the results, are available at https://github.com/saezlab/ligrec_decoupler.

### 4.4. Overlap Analysis

To compare the overlap between the interactions predicted by each method-resource combination, as a default we kept the 500 highest ranked interactions. We also considered the highest ranked 100, 250, and 1000 interactions for the CRC scRNA-Seq dataset. In case of ties, we considered the higher number of predicted interactions. We then generated a presence-absence matrix of predicted interactions with method-resource combinations. These matrices were subsequently used to calculate Jaccard indices and to cluster the results.

Activity per cell type was calculated using the highest ranked 500 hits for each method-resource combination. Cell type activity represents the proportion of interactions (or edges) that stem from or lead to a Source or Target cell type, respectively. In other words, a Source cell with a high cell type activity, in the broadest terms, can be inferred as an active ‘secretor of ligands’. We used the z-normalized average interaction rank for each possible combination of communicating cell types to estimate the cell pair ranks for each method-resource combinations. These patterns of pairwise communication activities we presented in a PCA plot. We created the heatmaps with pheatmap ^53^ (v1.0.12), using binary distances for the overlap heatmaps and euclidean distances for the other heatmaps.

Connectome, NATMI, and iTALK do not provide an explicit threshold to control for false positives and the highest ranking 500 hits were kept for each without any preceding filtering. For methods where a threshold was proposed by the authors, as in the case of CellChat, Squidpy, and SingleCellSignalR, we first filtered their results accordingly and the highest ranked interactions were obtained afterwards. Further, we used cluster-specific interaction measures for each method whenever available (Table 1).

The same analysis was also carried out using the cluster-unspecific measures from each method. The scaling done in Connectome (weight_scale) and NATMI (Specificity-based edge weight), and in particular the cluster label permutation of CellChat (p-values) and CellPhoneDB (p-values), explicitly reflect cell-cluster specific communication, thus we used their alternative measures. SingleCellSignalR and iTALK provide a single measure each and were hence excluded from this analysis.

### 4.5. Method Specifics

#### 4.5.1 CellChat

CellChat was run using default settings with 1000 permutations and the gene expression diffusion-based smoothing process was omitted. CellChat returned a number of significant interactions with p-values of 0 ranging from 221 to 12,208 depending on the resource (2,988 with its inbuilt resource), these made a considerable proportion of the significant hits (p-value <= 0.05), as they ranged from 237 to 12,971 (3,041 with its inbuilt resource). As such, because obtaining the highest ranked interactions based on p-values was infeasible, CellChat results were filtered according to p-values (p-value <= 0.05) and the highest probability scores were instead used in the method-resource analysis.

#### 4.5.2 Connectome

Connectome was run with its default settings using a Seurat object with processed gene expression counts. Results were filtered for differentially expressed genes (p-value <= 0.05), as identified via a Wilcoxon test, and Connectome’s scaled weights were used in the method-resource analysis.

#### 4.5.3 iTALK

iTALK was run with its default settings using the ‘DEG’ option which returns corrected p-values and logFold changes for each gene. Then transmitters and receivers with q-value <= 0.05 were kept. A differential expression product was calculated using z-scores of transmitters and receivers and subsequently used in the method-resource analysis.

#### 4.5.4 SingleCellSignalR

SingleCellSignalR was run with the processed gene counts, considering differentially expressed genes with a log2 fold change threshold of 1.5 or above. The highest LRscores which passed the recommended threshold of 0.5 were used in the method-resource comparison. The number of interactions predicted by SingleCellSignalR ranged between 159 to 7,240 (LRscore >= 0.5). The source code of SingleCellSignalR was modified to work with external resources (available at https://github.com/CostaLab/SingleCellSignalR_v1).

#### 4.5.5 NATMI

NATMI’s implementation is command-line based, thus a system command is invoked via R that calls the NATMI python module and passes the appropriate command line arguments. NATMI was run with its default settings using the processed gene expression matrix, converted from Seurat, and the specificity-based edge weights were used in the method-resource comparison. NATMI’s lrc2p resource was used as the default.

#### 4.5.6 Squidpy

Squidpy is called via reticulate ^59^ (https://rstudio.github.io/reticulate/) and the Seurat object is converted to [anndata](https://anndata.readthedocs.io/en/latest/index.html) ^60^ (https://anndata.readthedocs.io/) format in Python. The CellPhoneDB algorithm implementation was run via the Squidpy framework with 10,000 permutations, threshold of cells expressing transmitters and receivers of 0.1, and the minimum component expression was considered for complexes. For the method-resource comparison, we used the rank of p-values (p-value <= 0.05). Squidpy’s number of significant hits ranged between 60 to 2,927 depending on the resource.

## 5. Acknowledgements

This work was supported in part by the European Union’s Horizon 2020 research and innovation program (860329 Marie-Curie ITN “STRATEGY-CKD”) and the German Federal Ministry of Education and Research (Bundesministerium für Bildung und Forschung BMBF) Computational Life Sciences LaMarck grant no. 031L0181B), awarded to JSR. This work was in part funded by the clinical research unit CRU344 supported by the German Research Foundation (DFG) and the E:MED Consortia Fibromap funded by the German Ministry of Education and Science (BMBF) awarded to I.C.

We express our gratitude to Erick Armingol, Pau Badia i Mompel, Hratch Baghdassarian, Luz Garcia-Alonso and Suoqin Jin for their helpful feedback and discussions.

## 7. Authors contributions

JSR conceived the project. DD set up the framework used in the comparison of method-resource combinations, with the help of JSN and DT. JSN was supervised by IC. DT and CB created the resource analysis plots and pipeline, later extended by DD. JSR supervised the project with the help of AV and AD. DD wrote the manuscript together with AV, CB, DT, and JSR. HK, RORF, and BS performed preliminary and supplementary analyses that helped shape the work presented here and the future directions of this project. All authors contributed to the ideas presented in this work and all revised the final version of the manuscript.

## Supplementary Materials

**Supplementary Figure S1.**
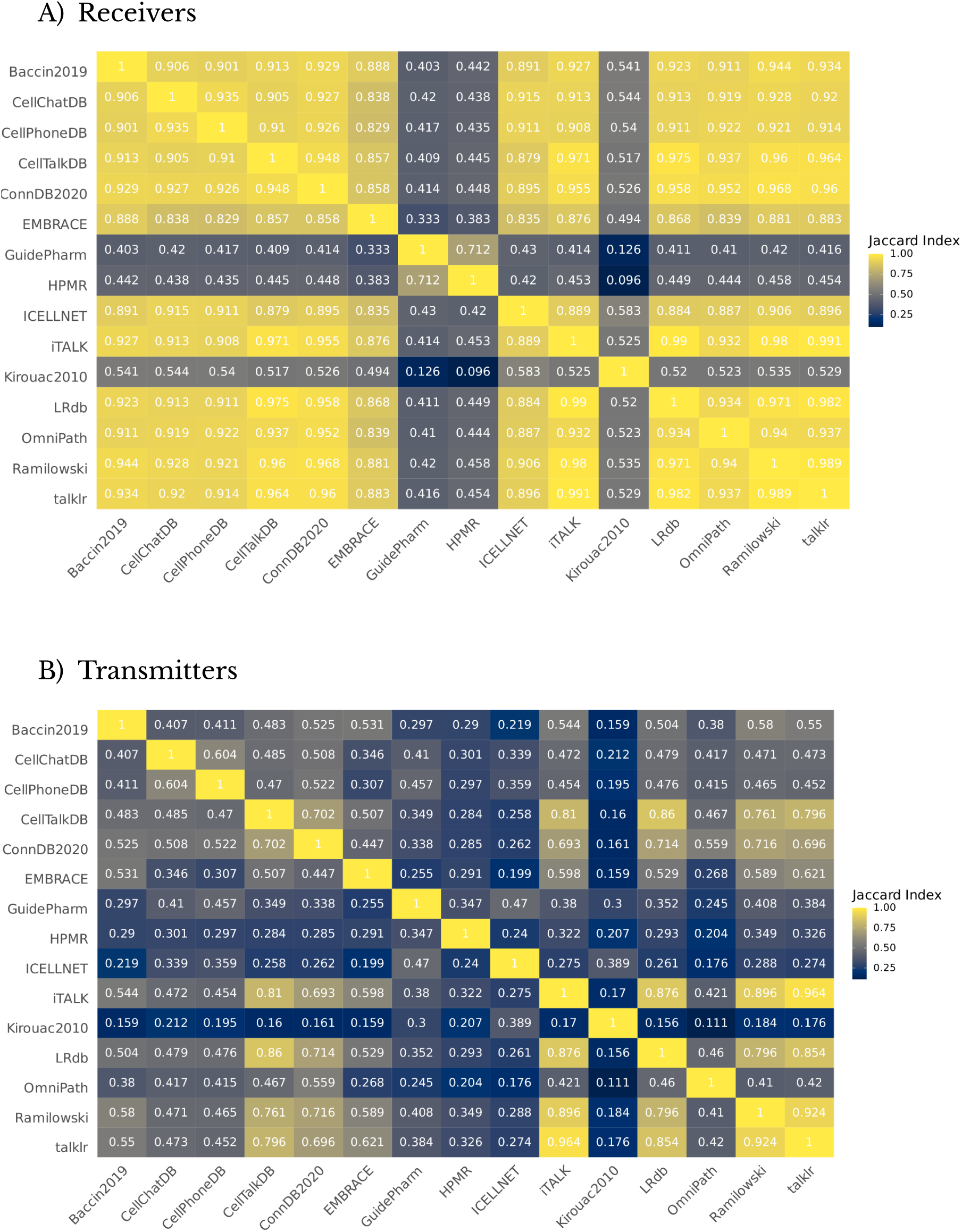
Jaccard Indices of A) Receivers and B) Transmitters from different resources.

**Supplementary Figure S2.**
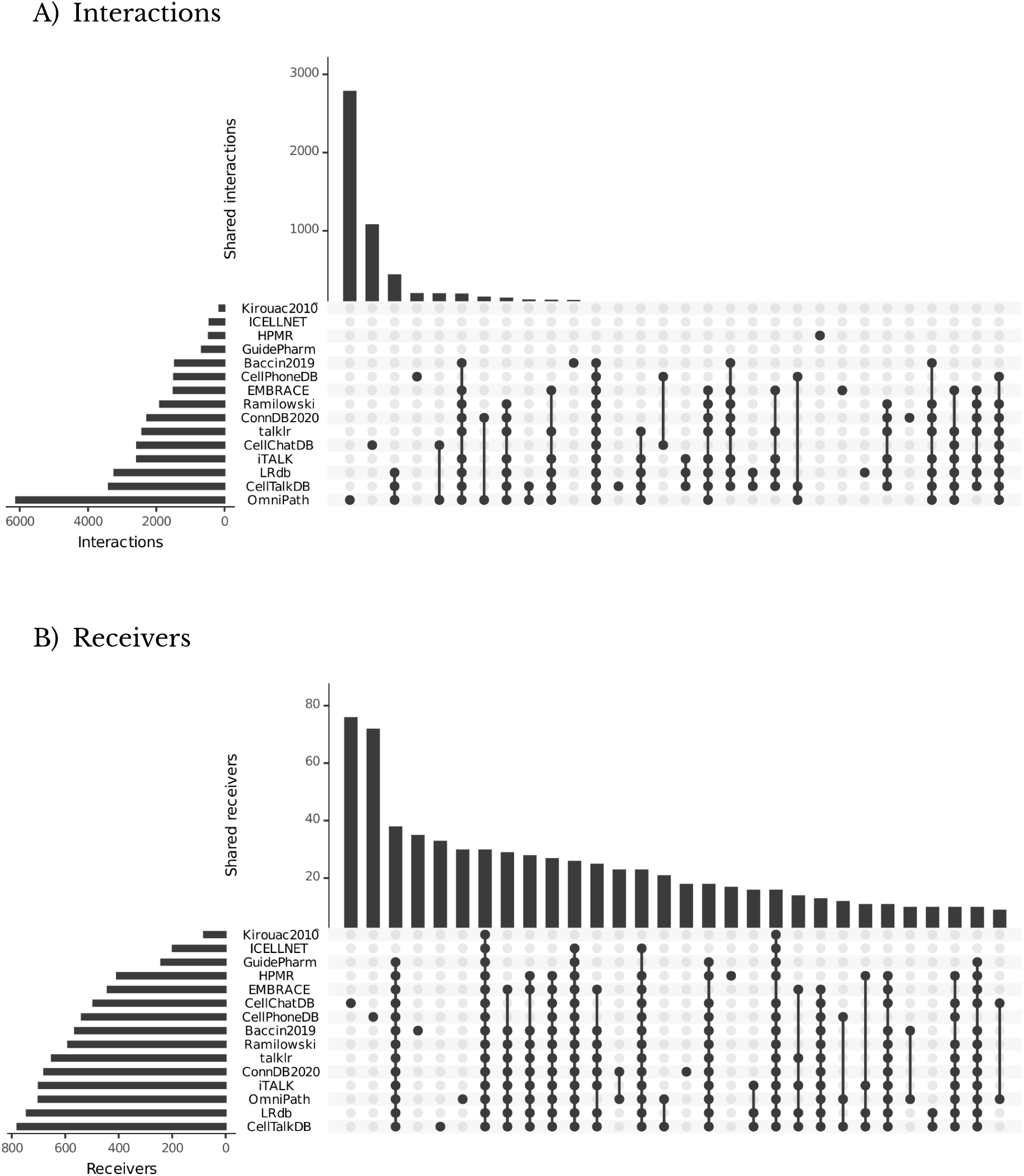
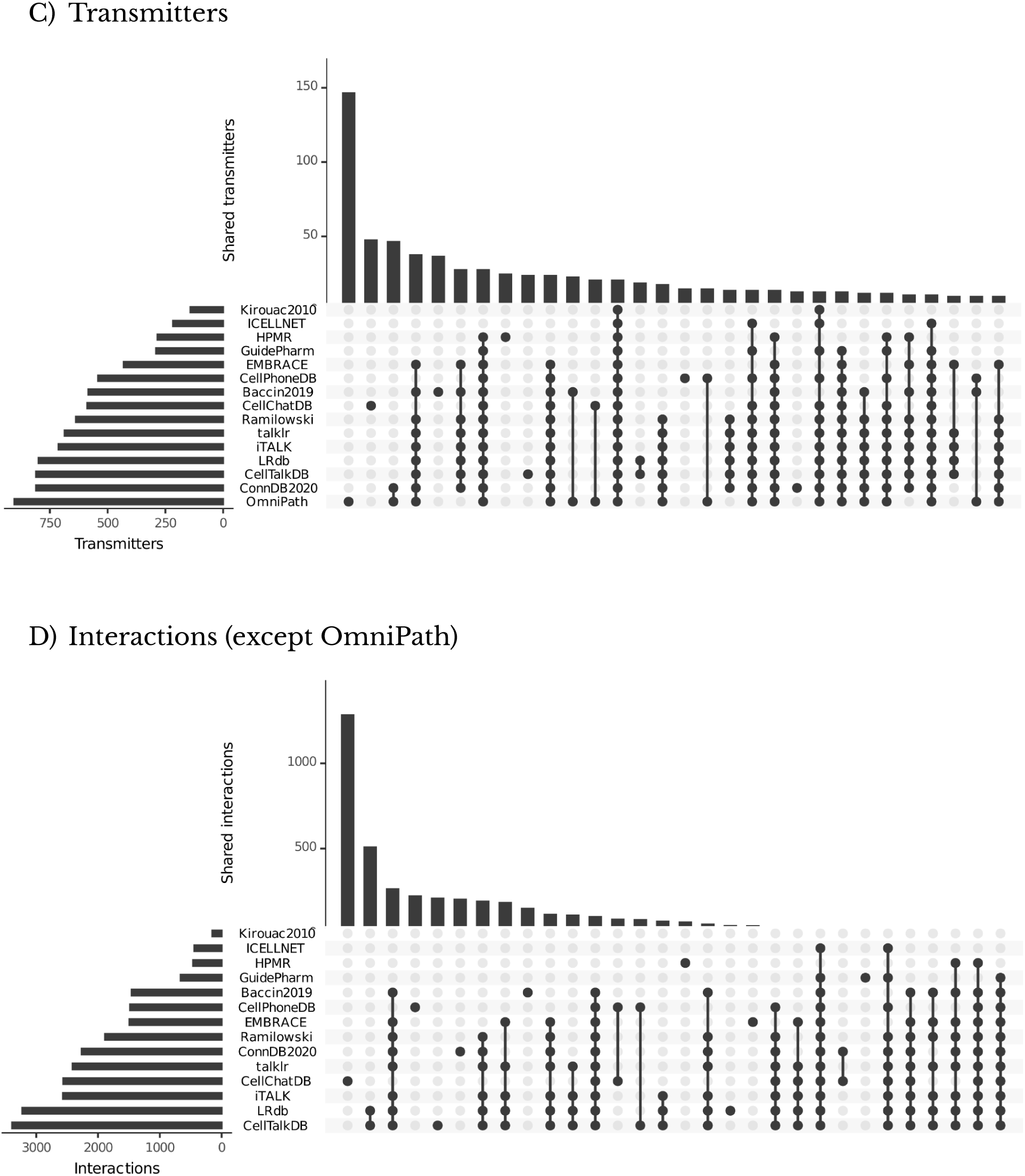
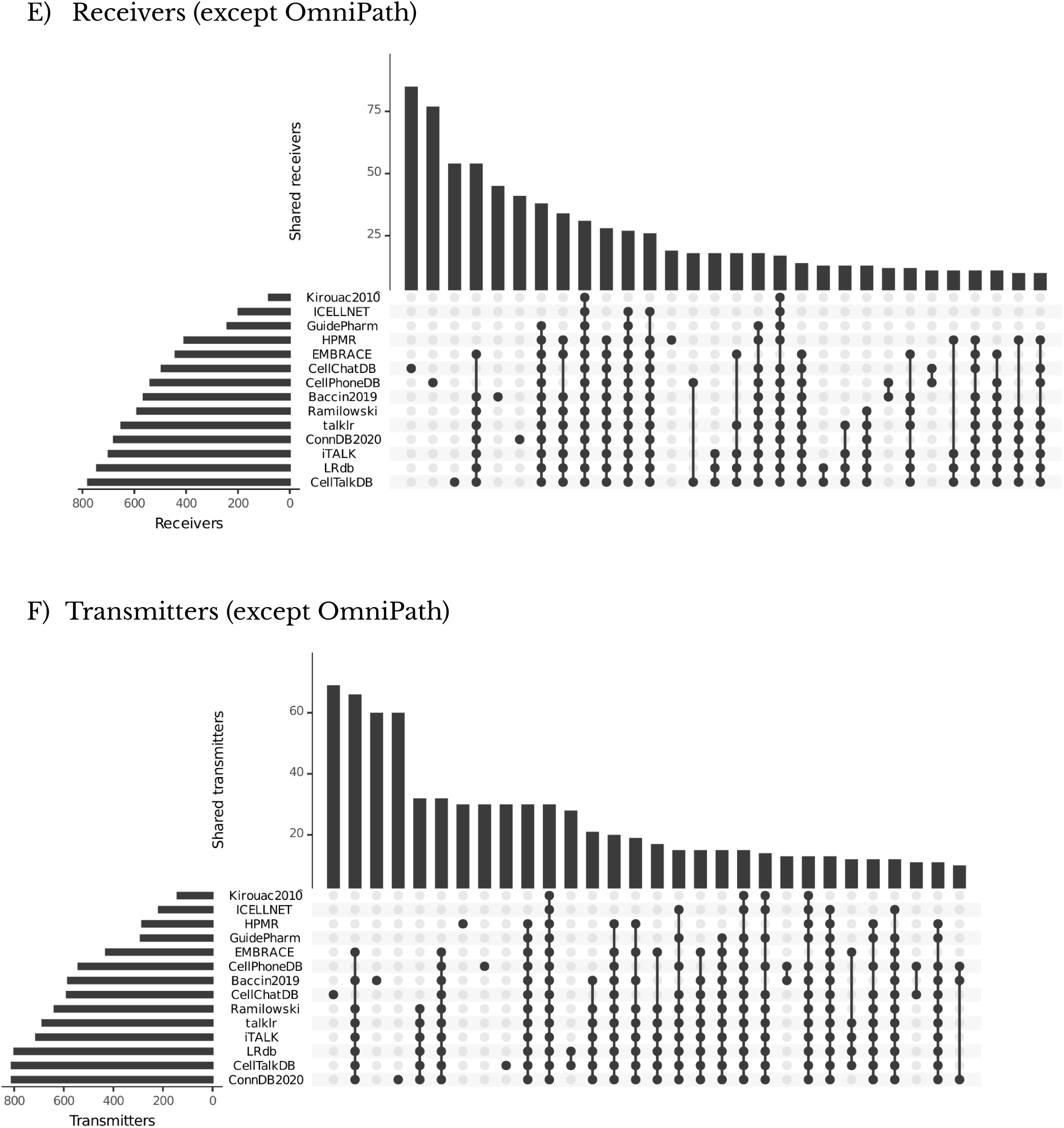
Upset plots representing the shared Interactions, Receivers, and Transmitters between all resources (A-C) and all resources except OmniPath (D-F).

**Supplementary Figure S3.**
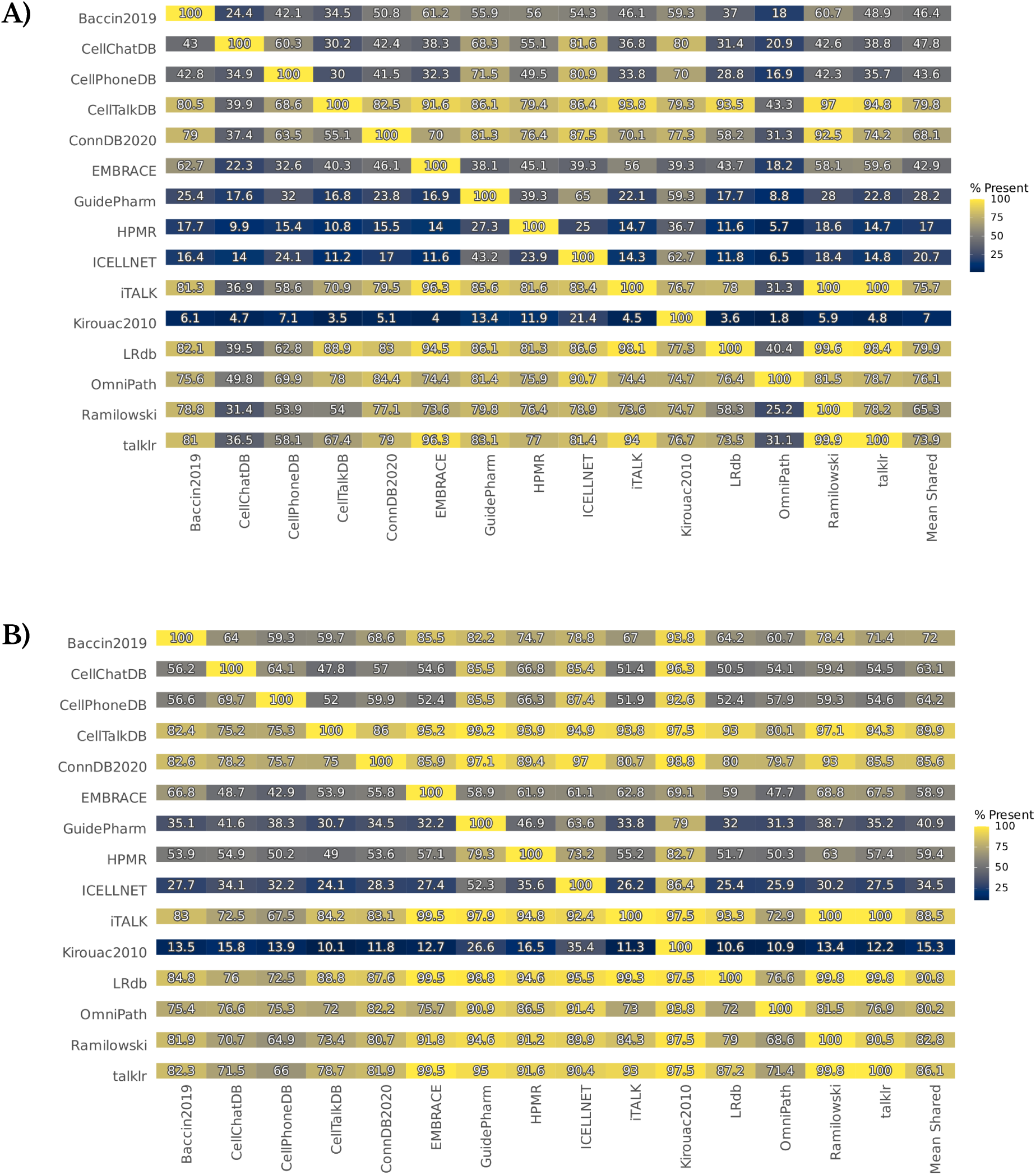
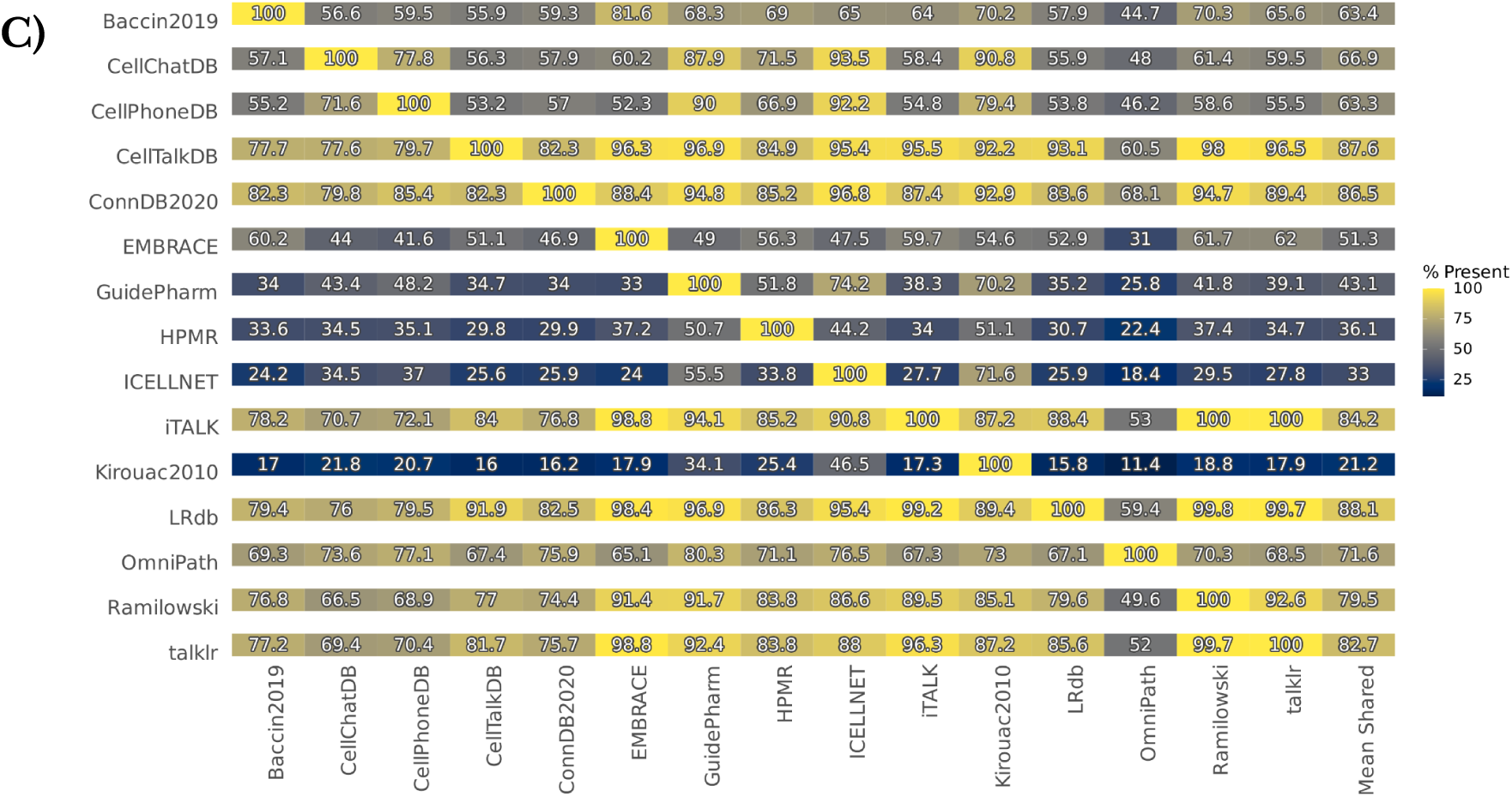
A) Interactions B) Receivers and C) Transmitters present in each resource when taken from the rest of the resources. Note these plots are asymmetric and represent the % of interactions from the resources on the X axis found in each resource on the Y axis.

**Supplementary Figure S4.**
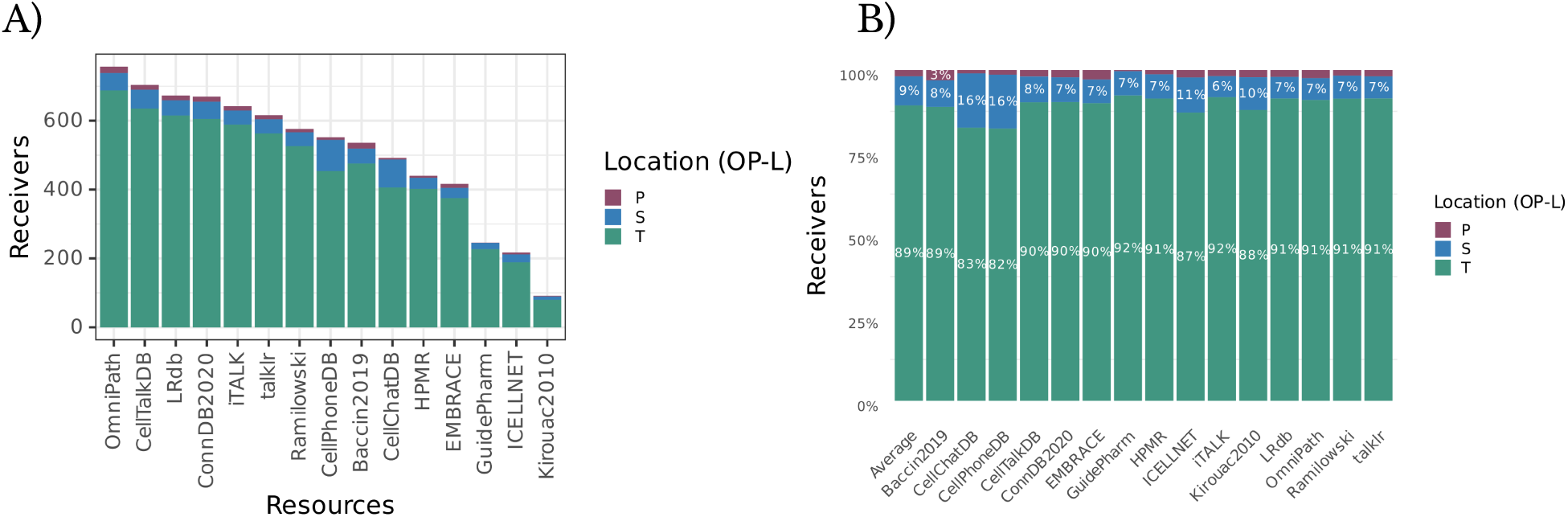
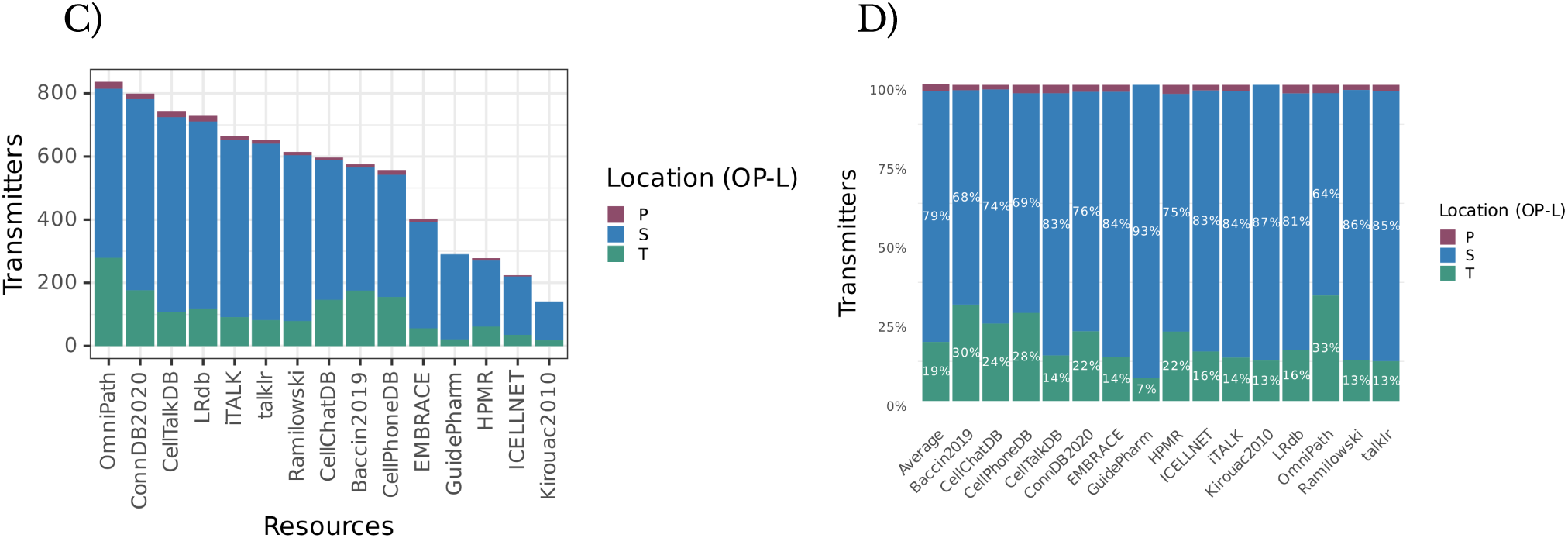
Numbers and Percentages of Subcellular locations annotations of Receivers (A-B) and Transmitters (C-D) for each CCC resource. S, P and T stand for Secreted, Peripheral plasma membrane, and Transmembrane plasma membrane proteins, respectively.

**Supplementary Figure S5.**
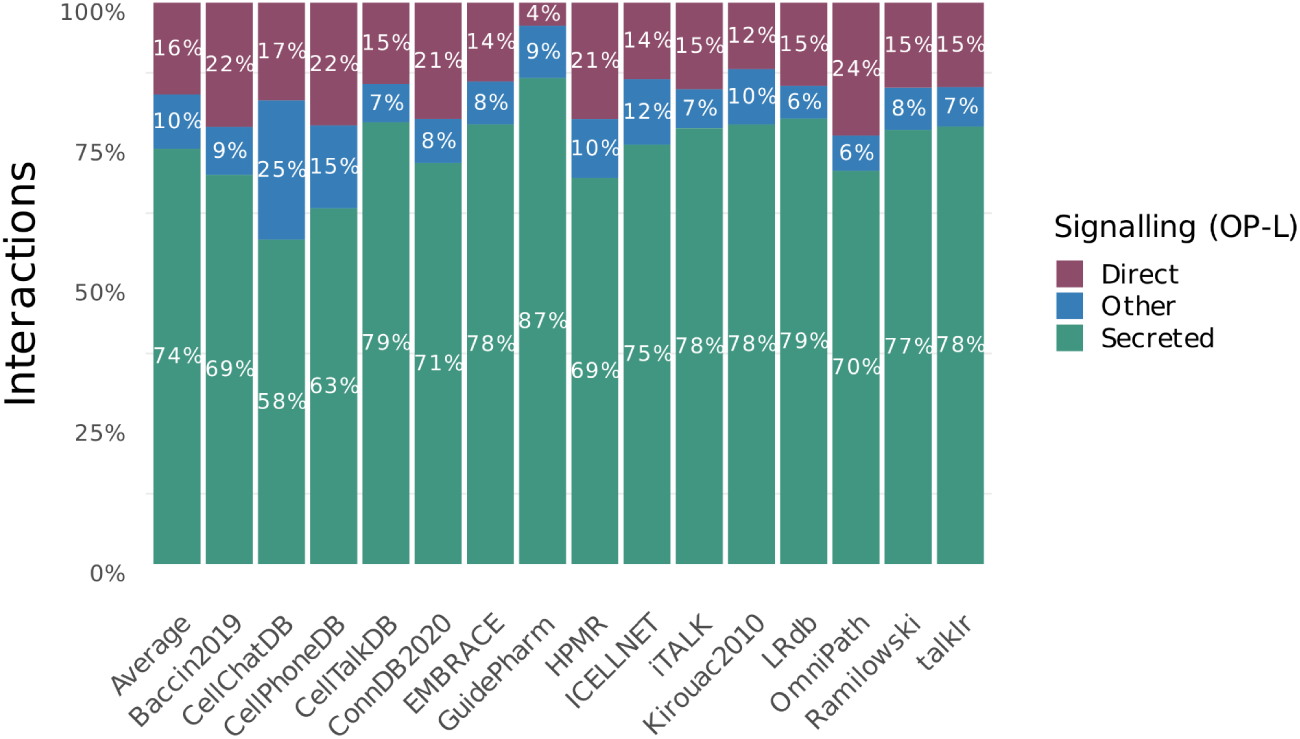
Percentages per Signalling category according to OmniPath locations (OP-L) for each CCC resource.

**Supplementary Figure S6.**
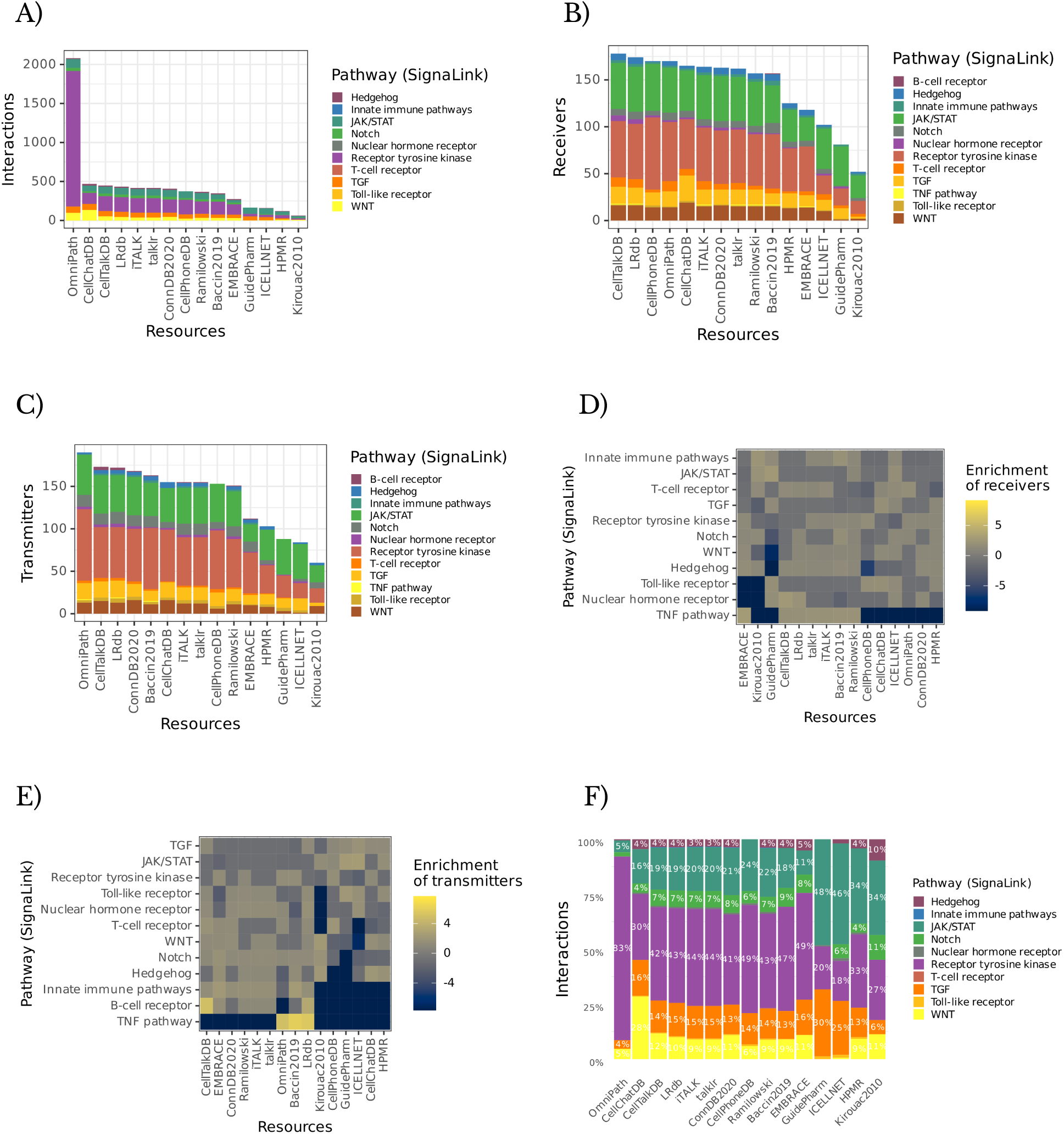
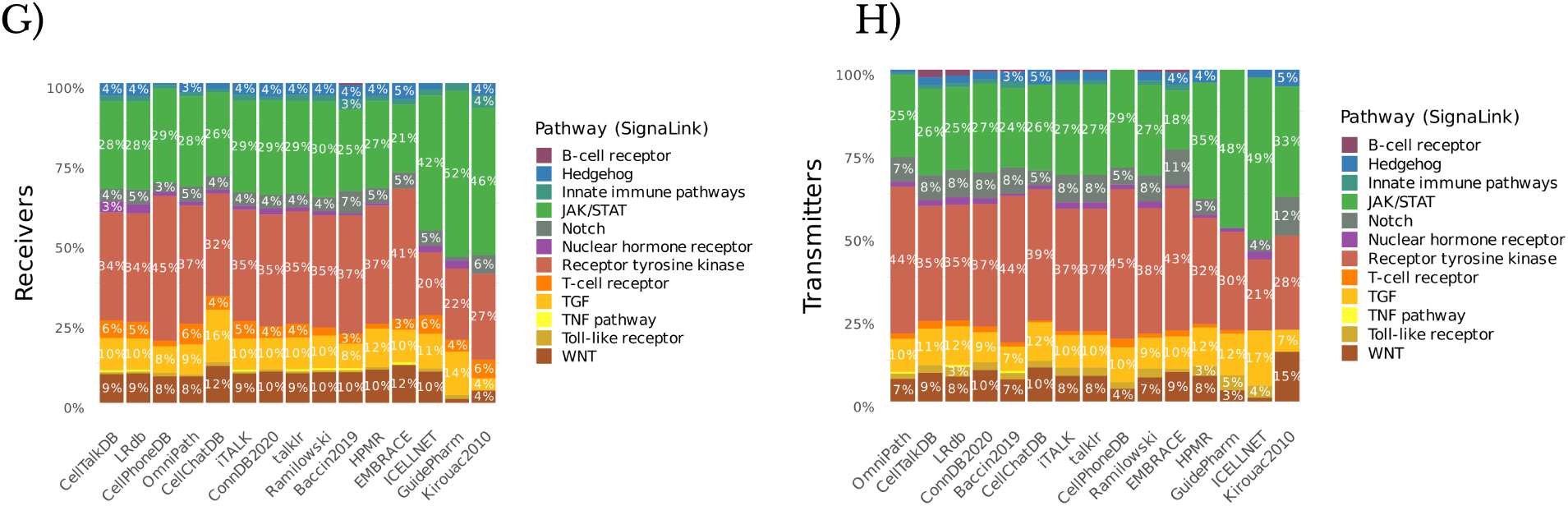
Number of matches to A) Interactions, B) Receivers and C) Transmitters, Enrichment Scores for their Receivers and Transmitters (D-E), and the Percentages of Interactions, Receivers and Transmitters (F-H) matched to the SignaLink database per resource.

**Supplementary Figure S7.**
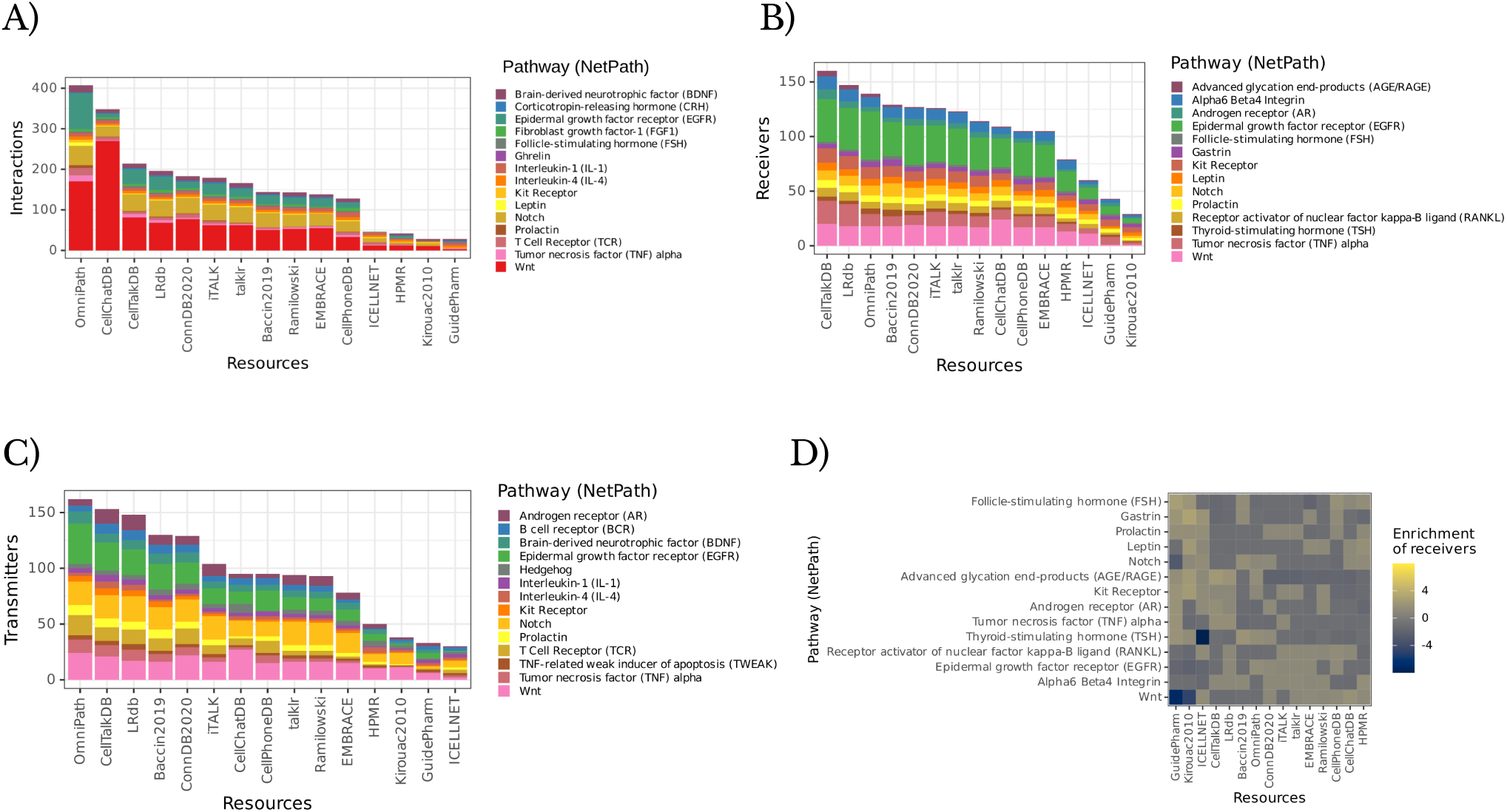
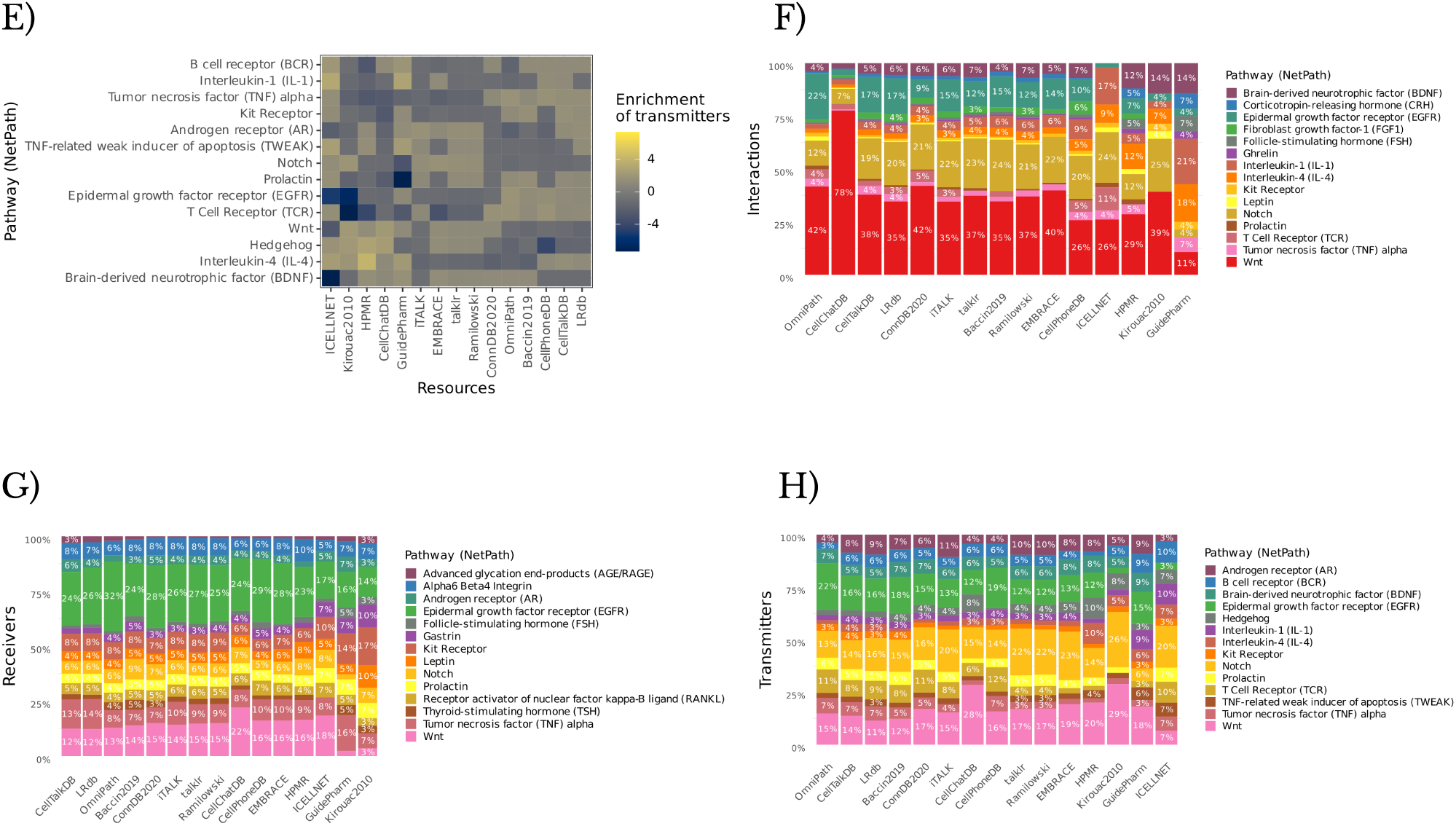
Number of matches to A) Interactions, B) Receivers and C) Transmitters, Enrichment Scores for their Receivers and Transmitters (D-E), and the Percentages of Interactions, Receivers and Transmitters (F-H) matched to the NetPath database per resource.

**Supplementary Figure S8.**
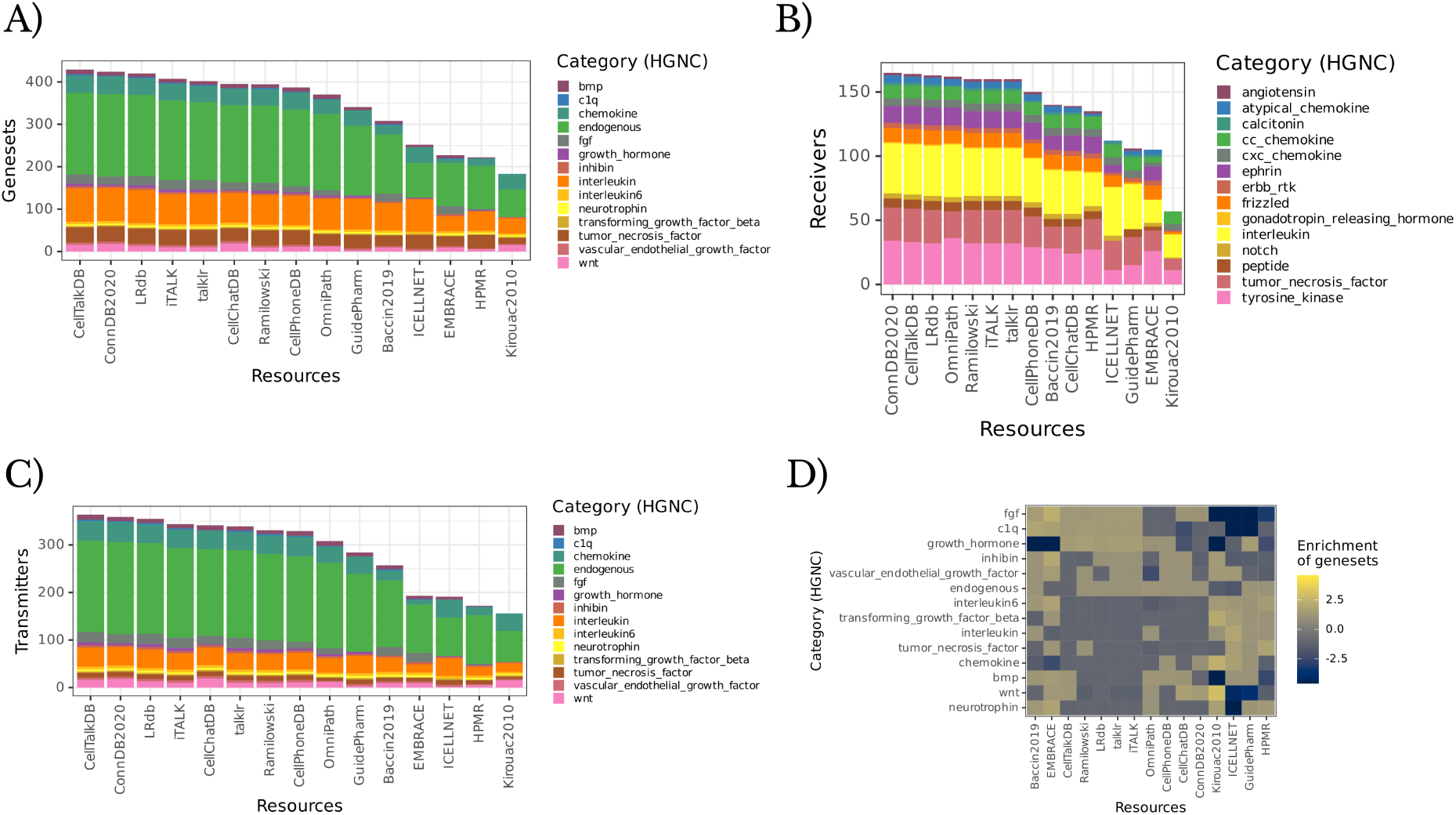
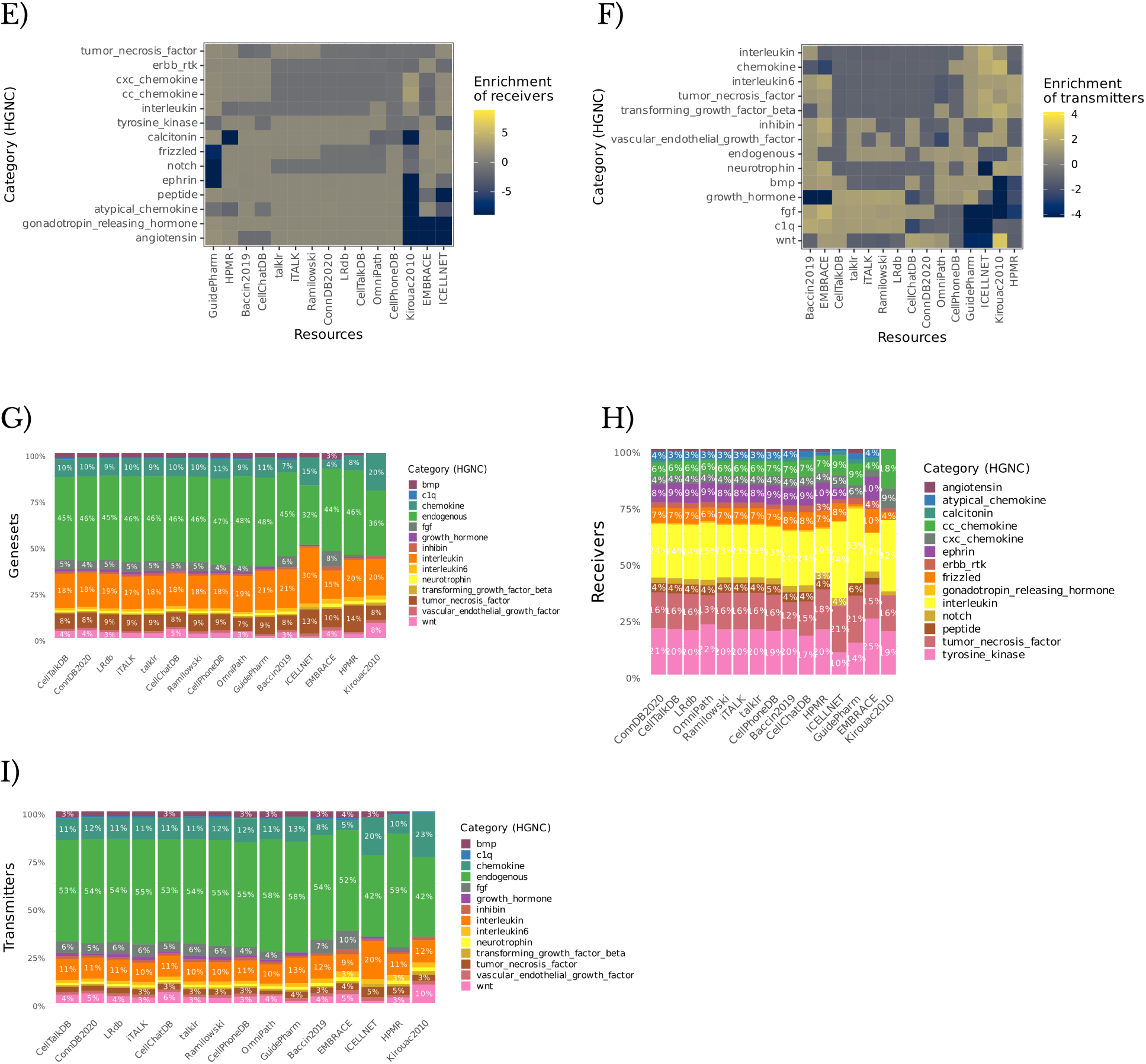
Number of matches to A) Merged Sets of Receivers and Transmitters, B) Receivers and C) Transmitters, their corresponding Enrichment Scores (D-F), and Percentages (G-I) per resource matched to the HGNC database.

**Supplementary Figure S9.**
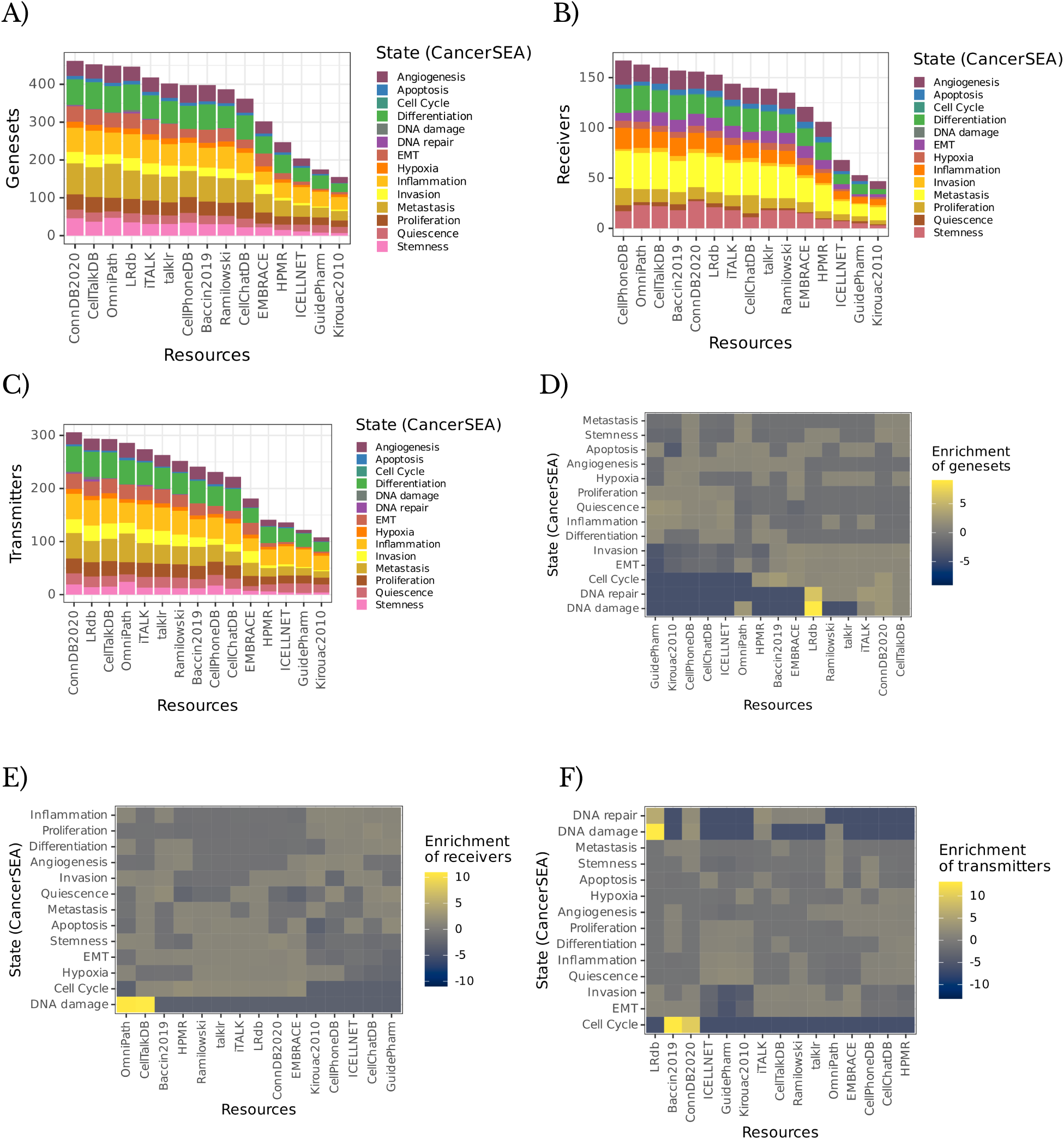
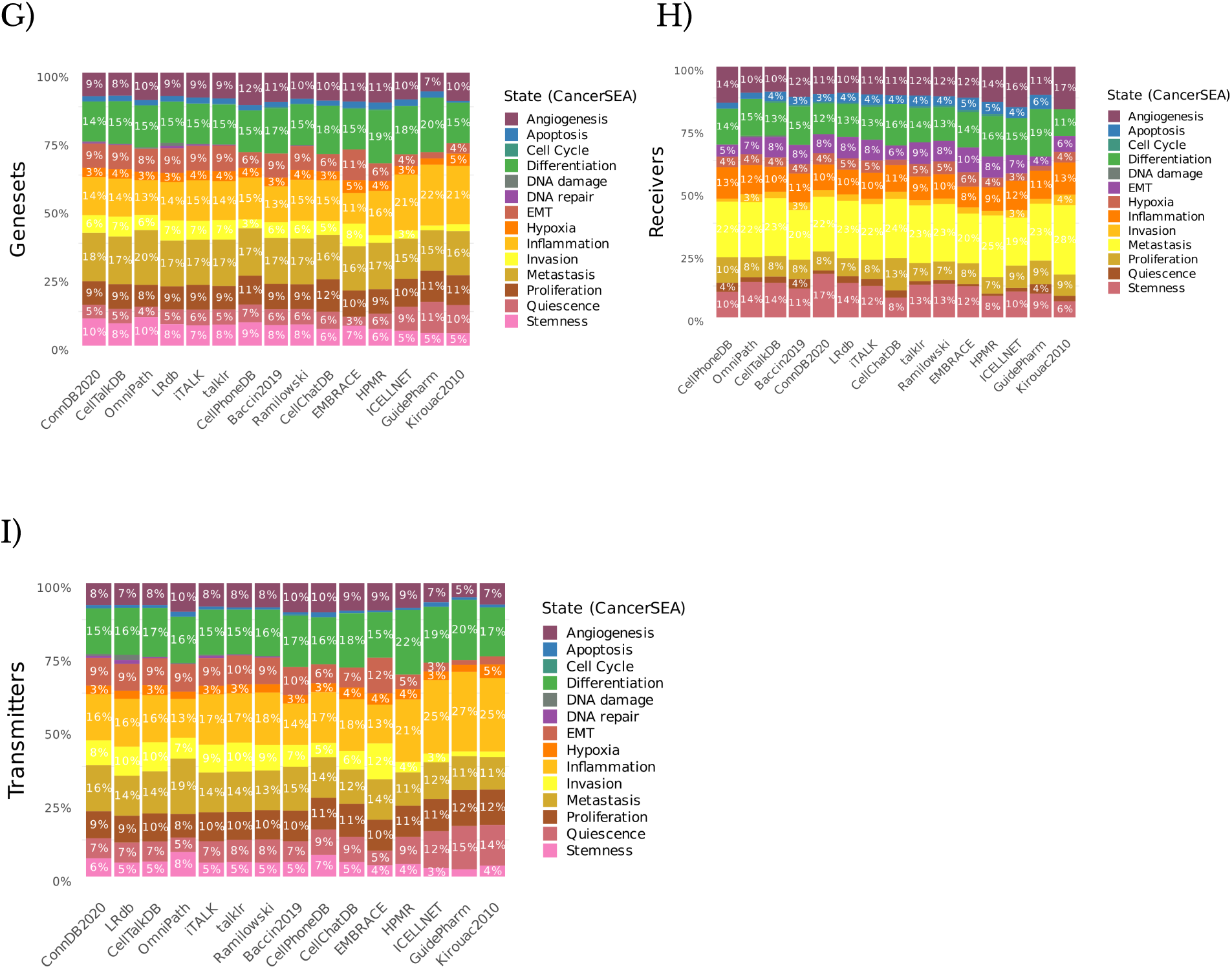
Number of matches to A) Merged Sets of Receivers and Transmitters, B) Receivers and C) Transmitters and their corresponding (D-F) Enrichment Scores, and Percentages (G-I) per resource matched to the CancerSEA database.

**Supplementary Figure S10.**
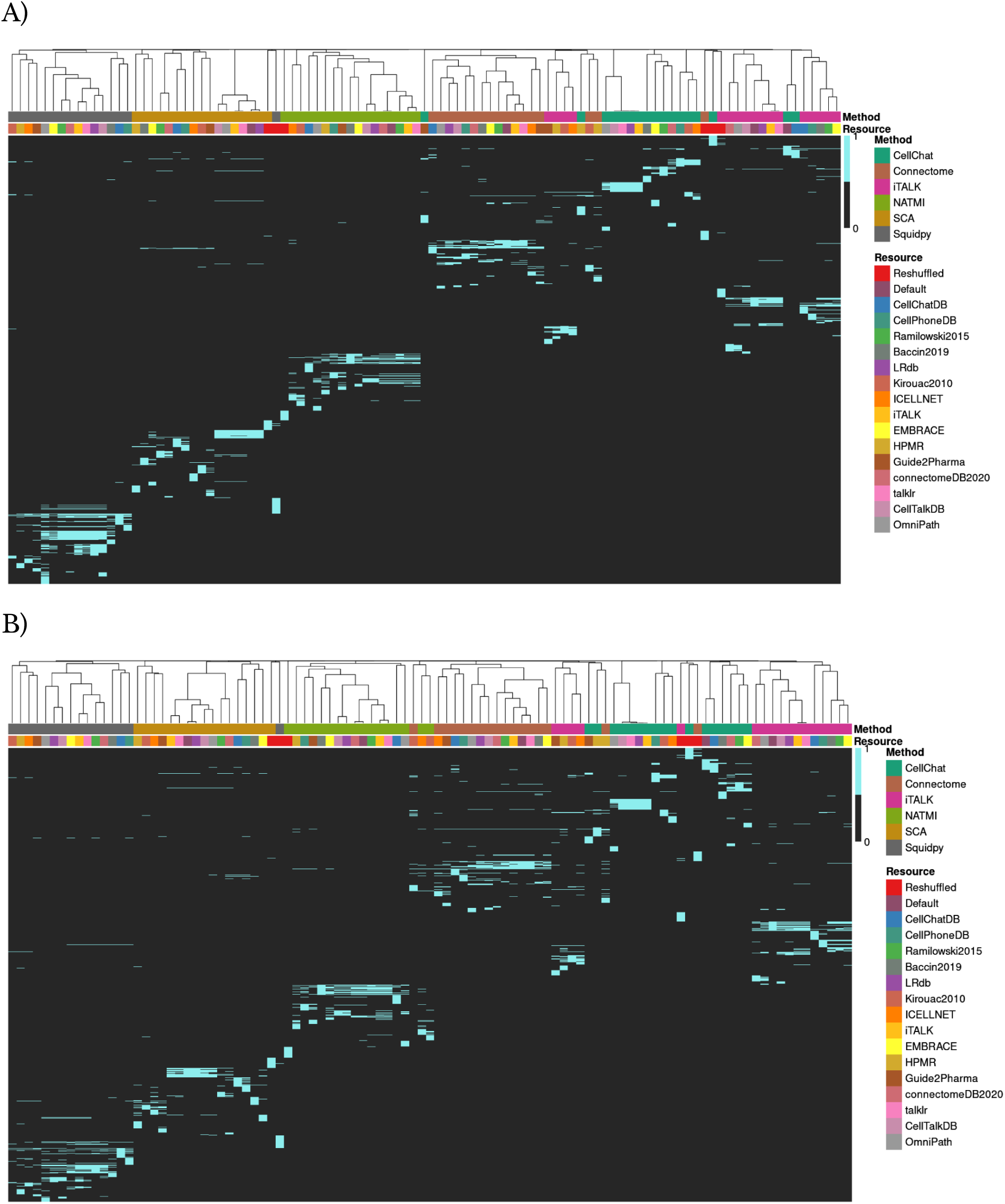
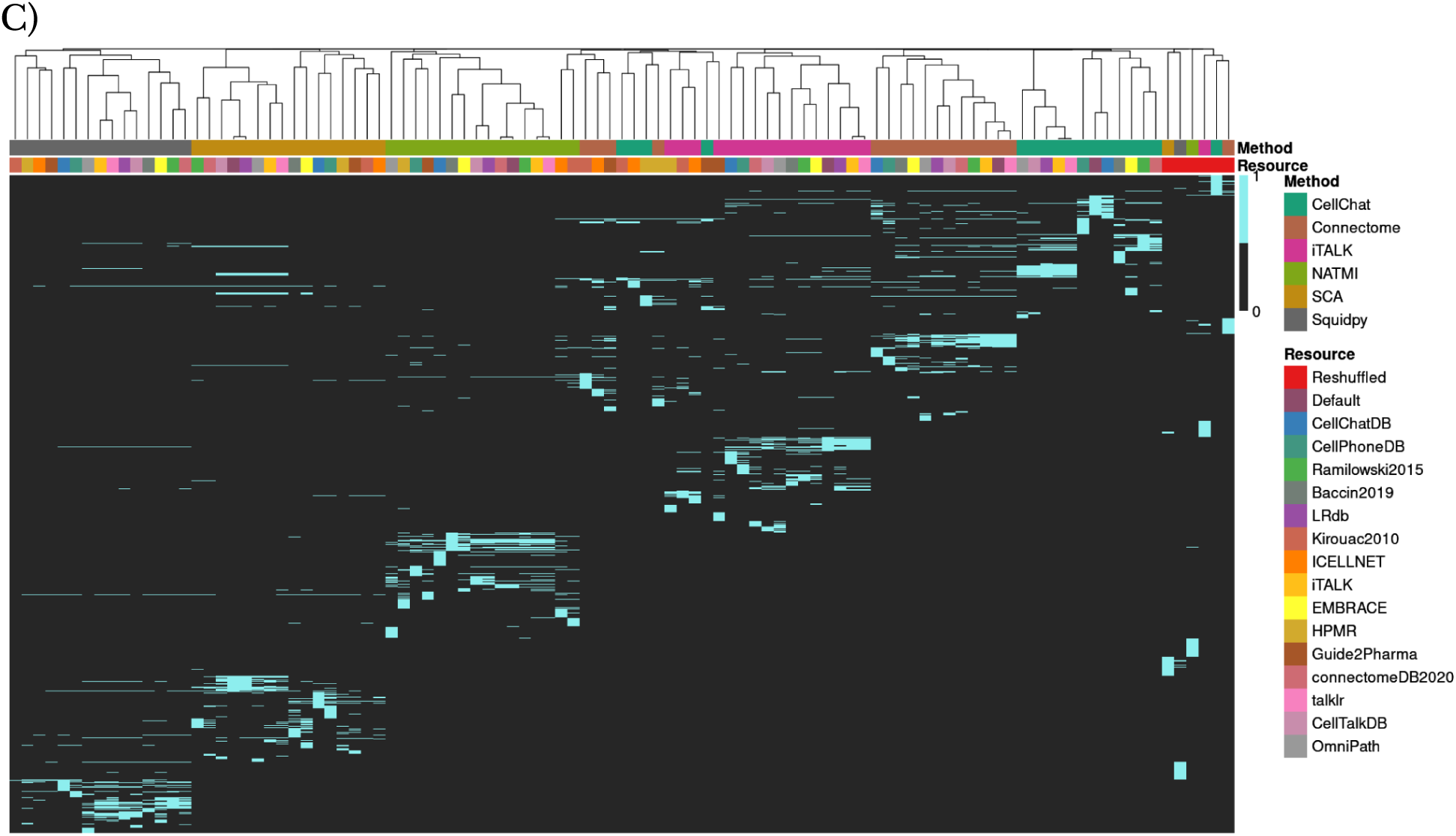
Overlap in the A) 100, B) 250, and C) 1000 highest ranked CCC interactions between different combinations of methods and resources. Method-resource combinations were clustered according to binary (Jaccard index) distances. SCA refers to the SingleCellSignalR method.

**Supplementary Figure S11.**
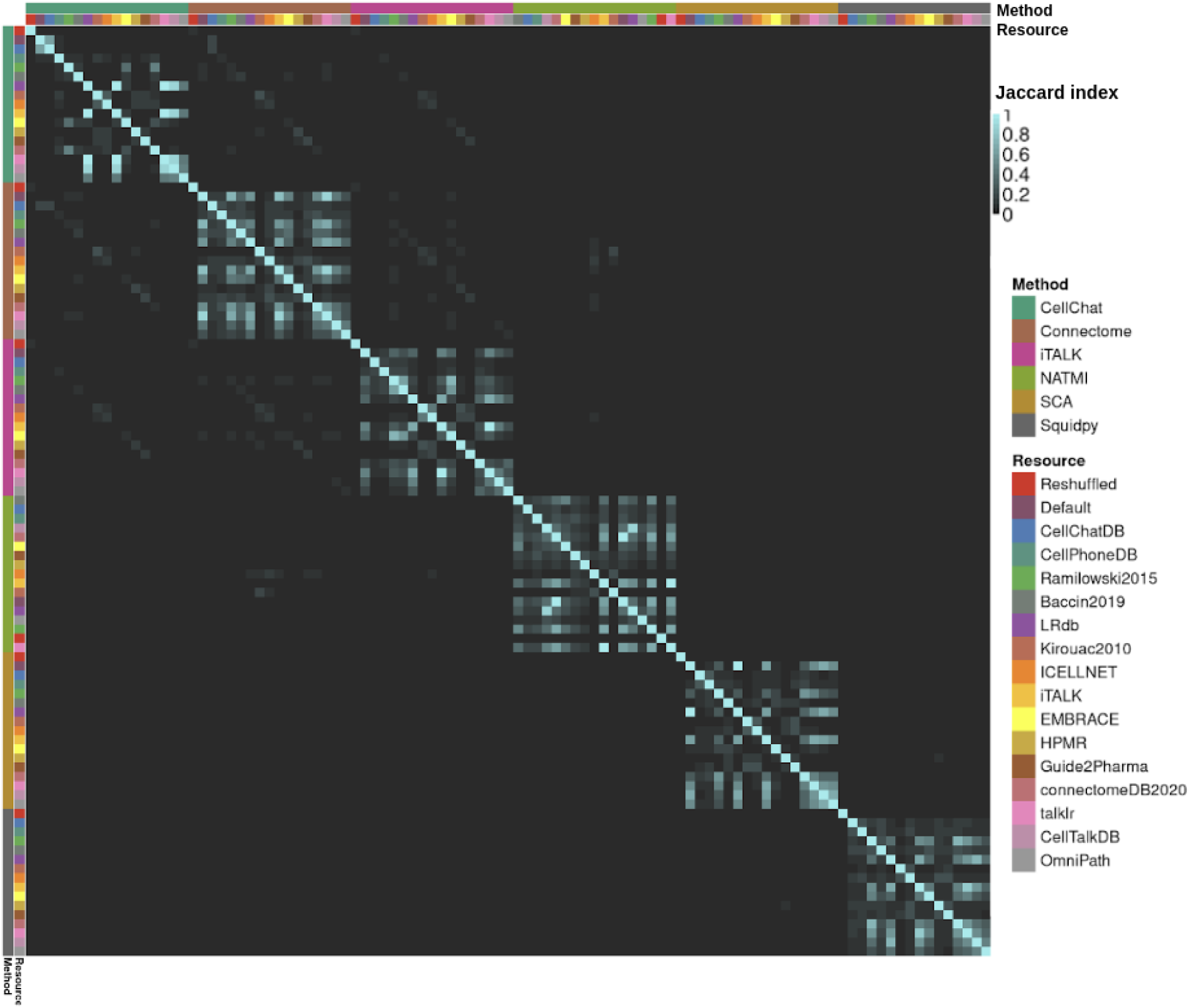
Jaccard indices for the 500 highest ranked interactions obtained from each method-resource combination.

**Supplementary Figure S12.**
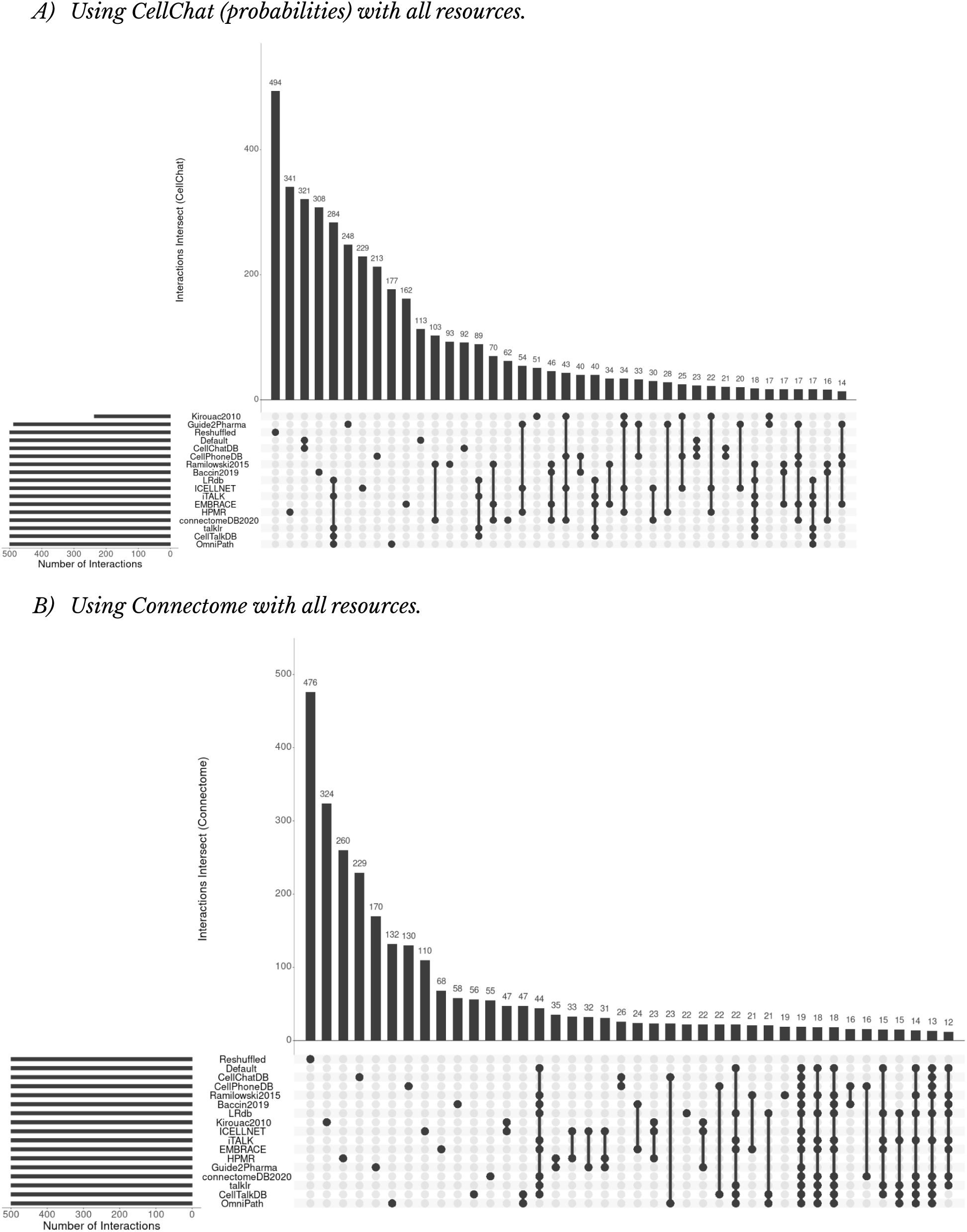
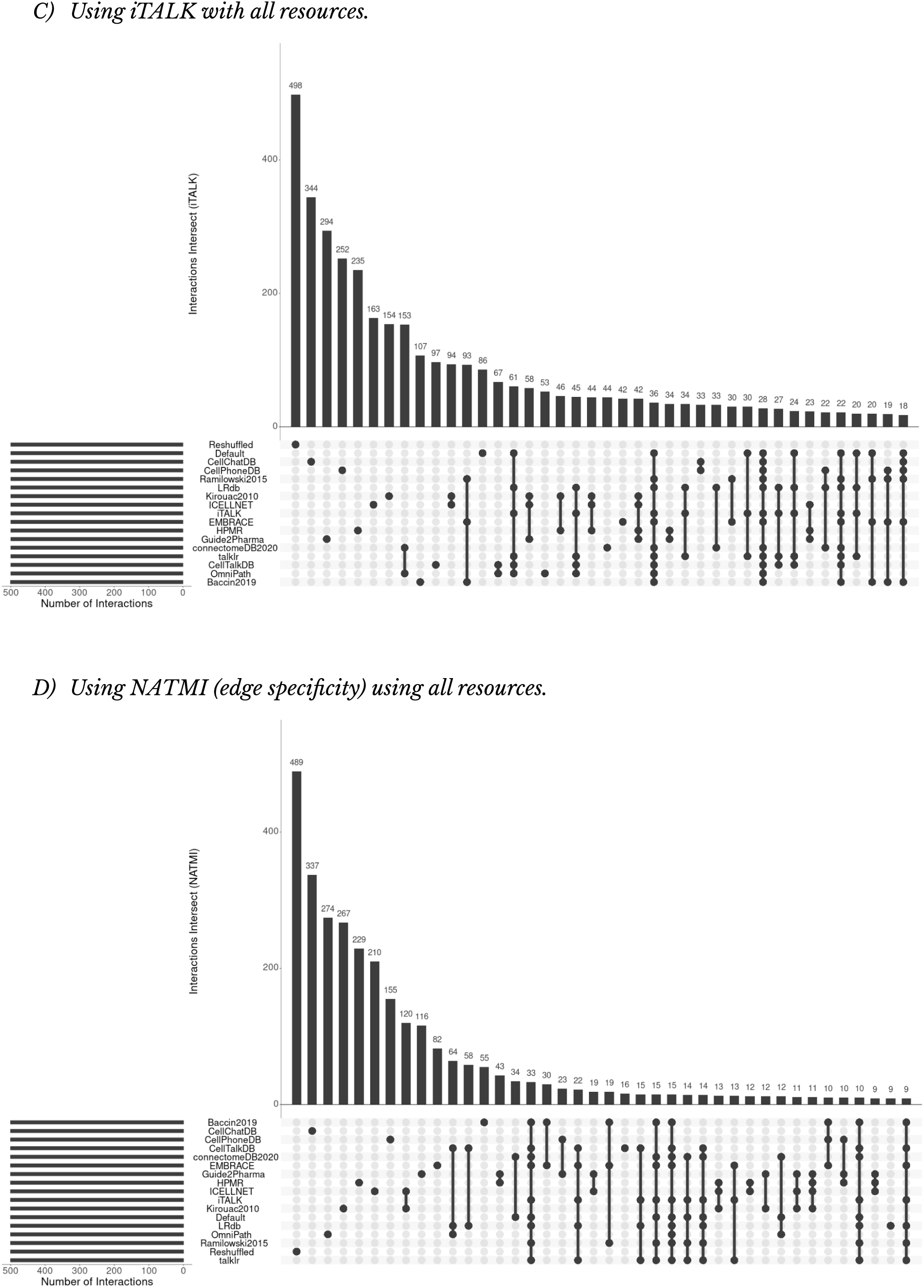
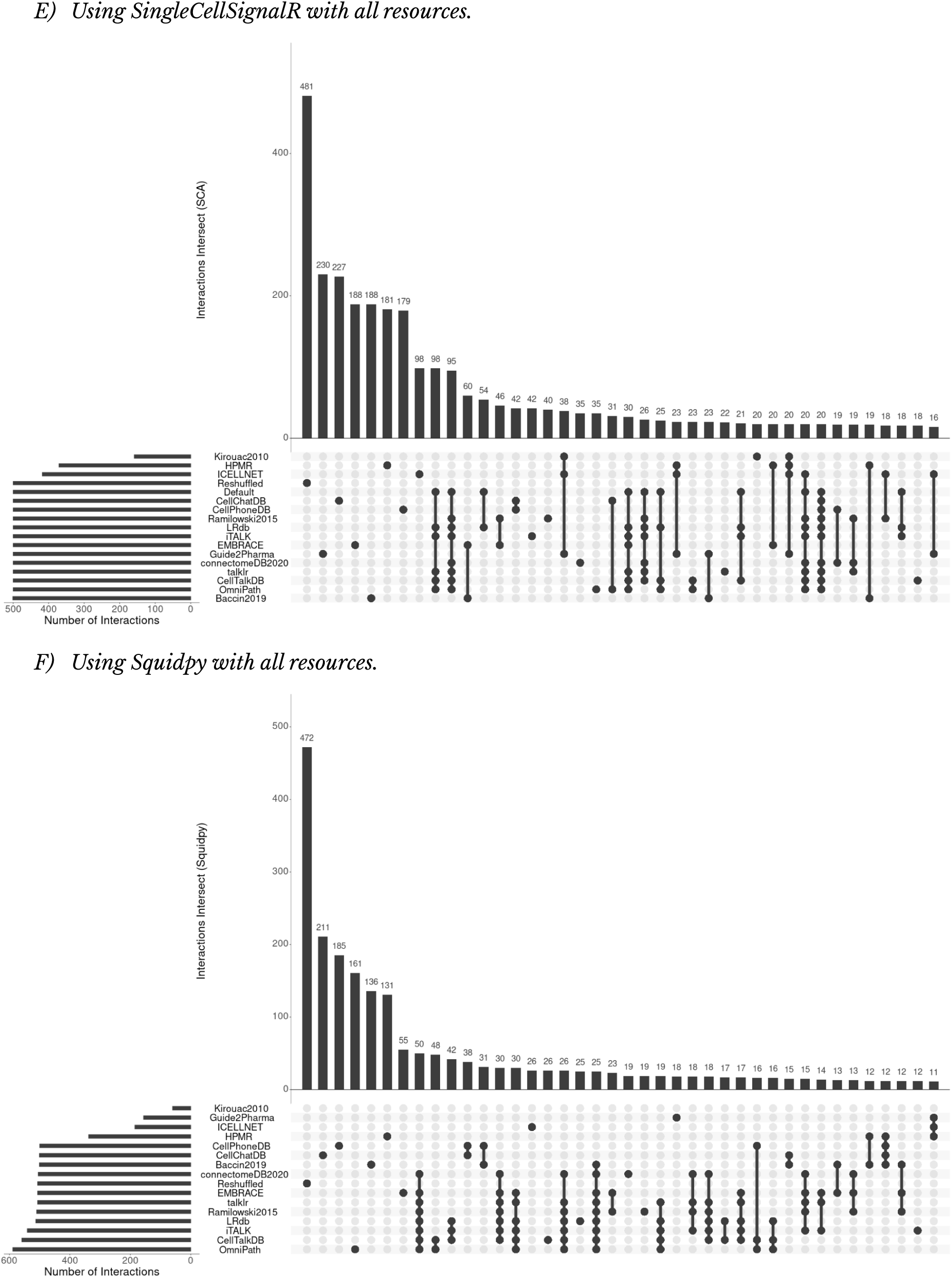
Upset plot showing the overlap between the 500 highest ranked interactions using the same method with all resources.

**Supplementary Figure S13.**
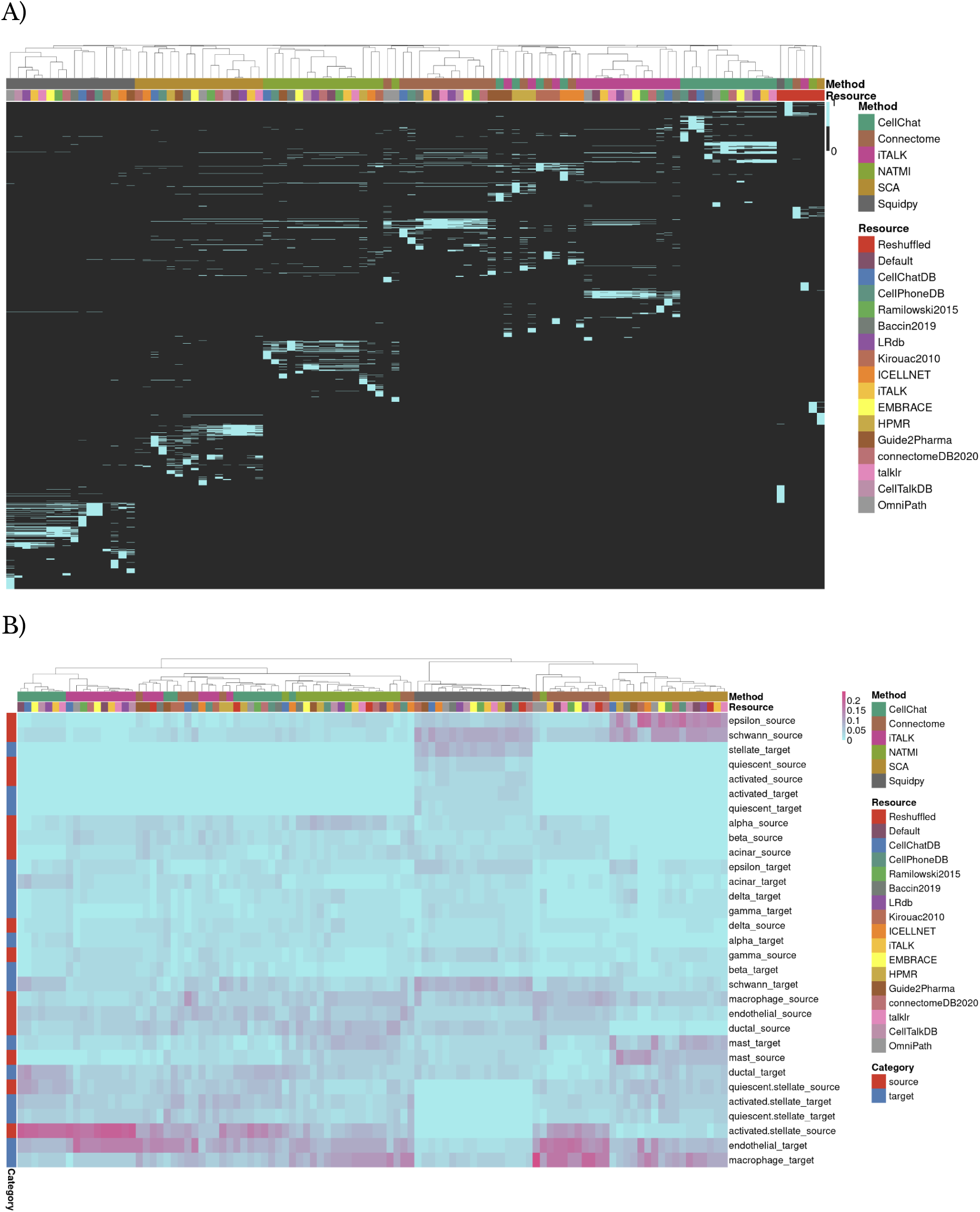
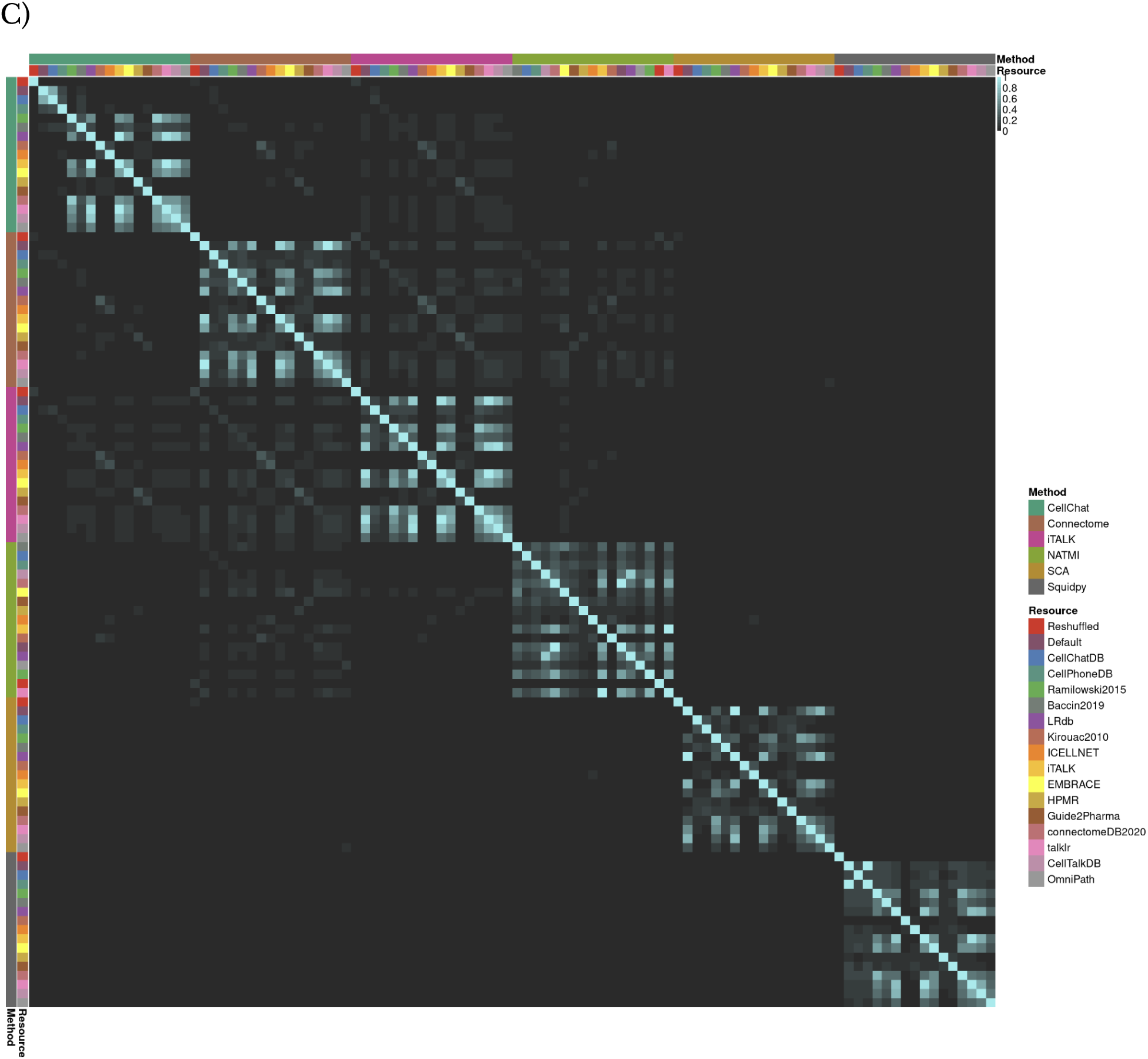
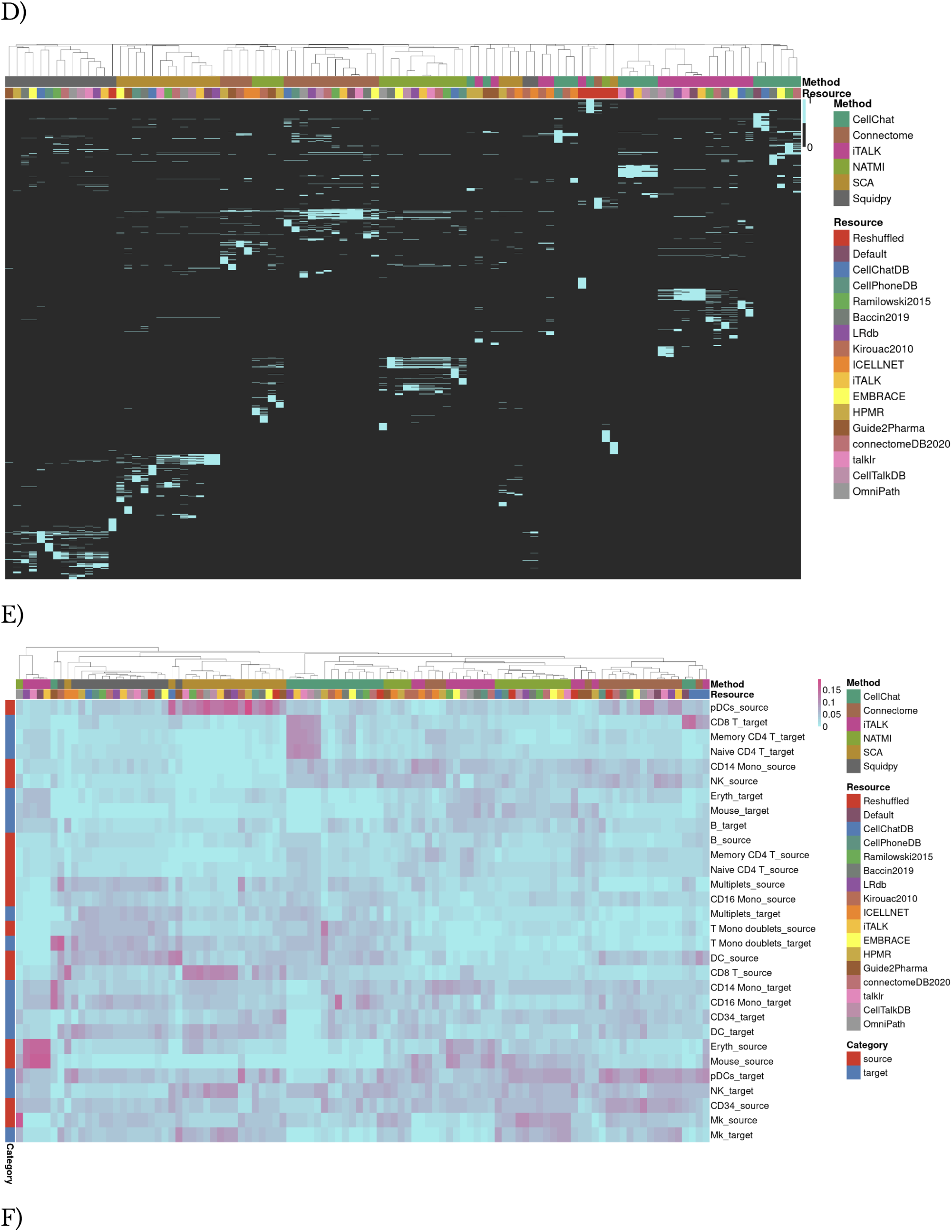
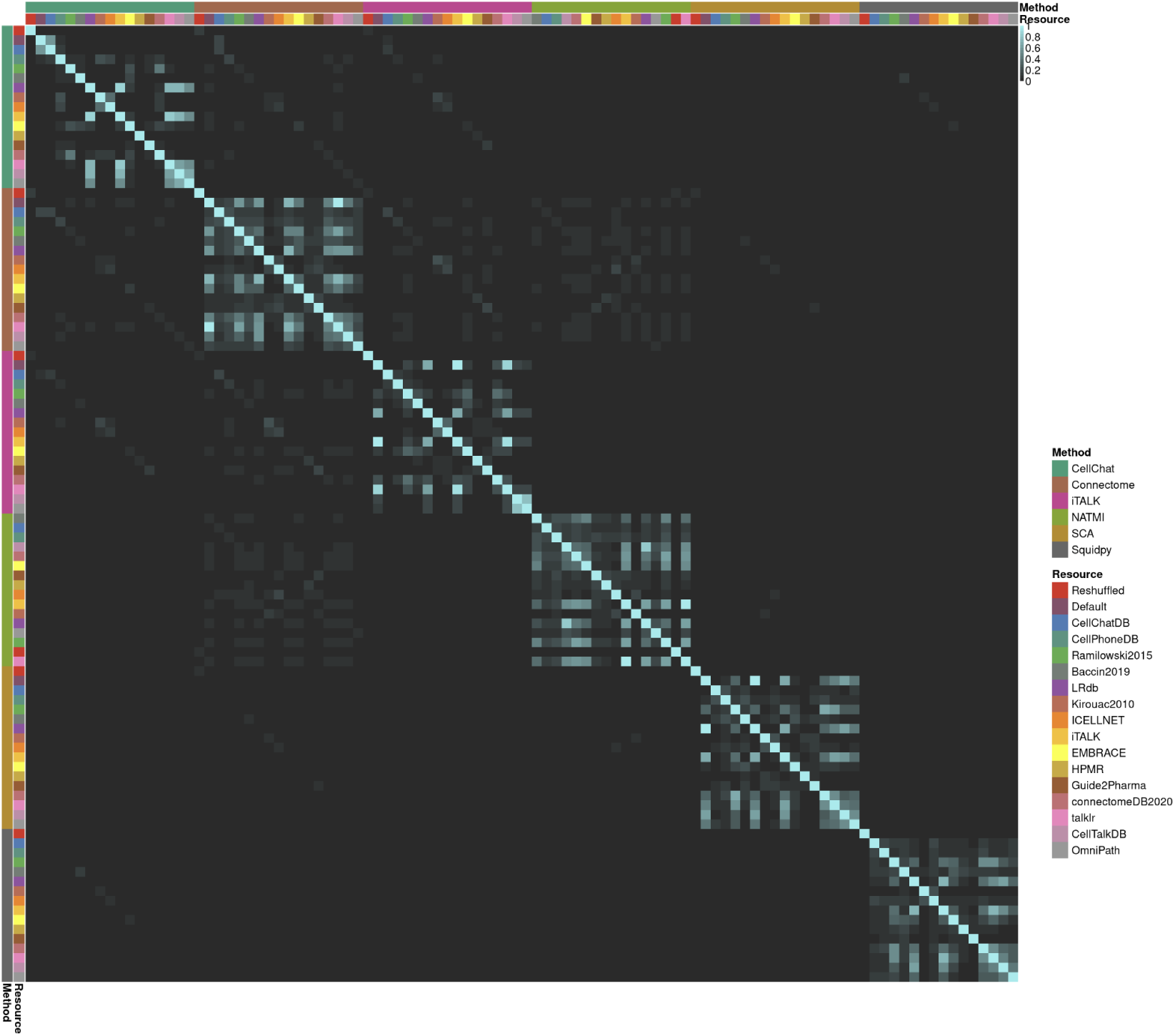
Overlap, Jaccard indices, and Activity per Cell type in the 500 highest ranked interactions between different combinations of methods and resources for Pancreatic islet (A-C) and Cord Blood Mononuclear Cells (D-F) scRNA-Seq datasets, respectively.

**Supplementary Figure S14.**
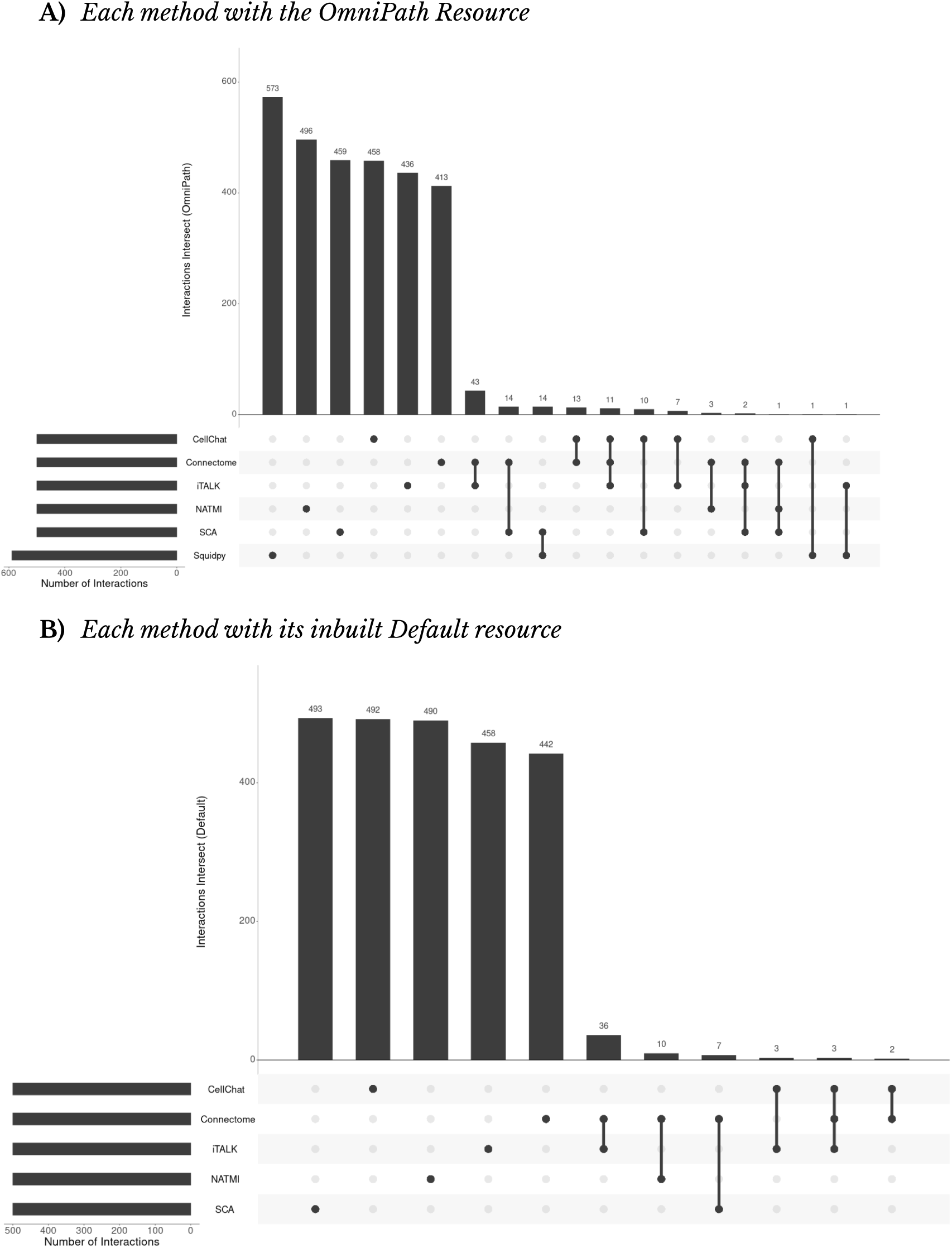
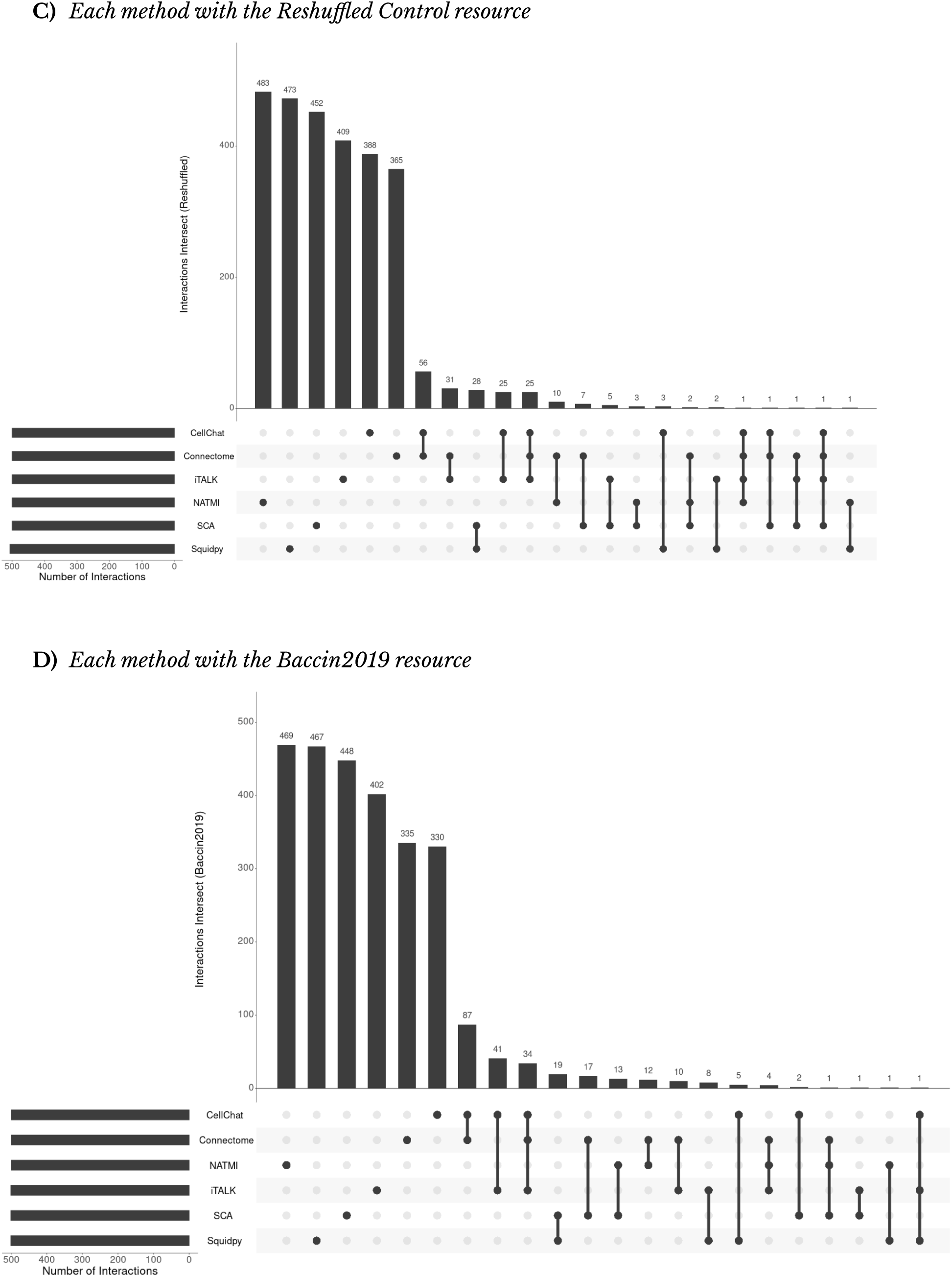
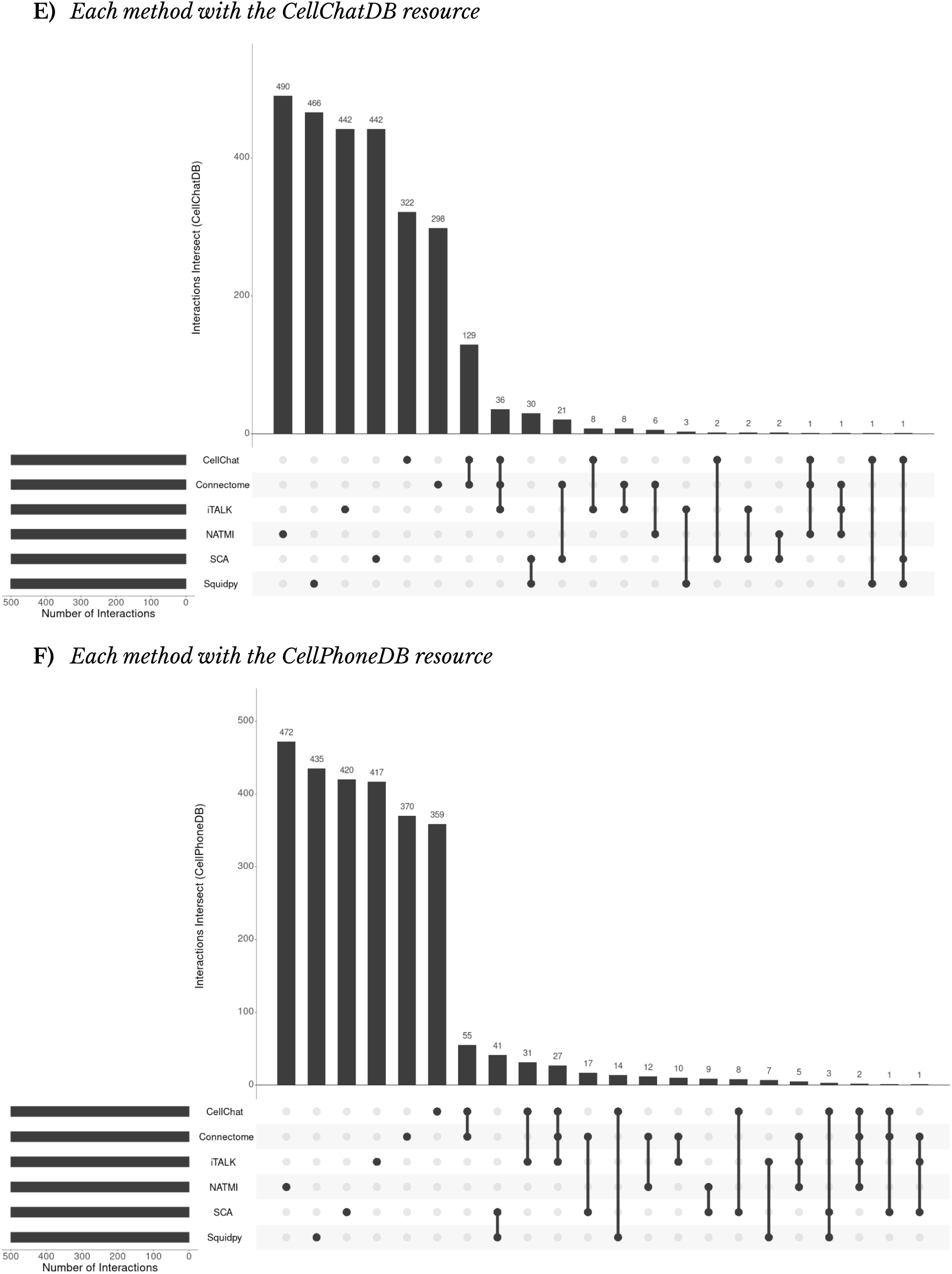
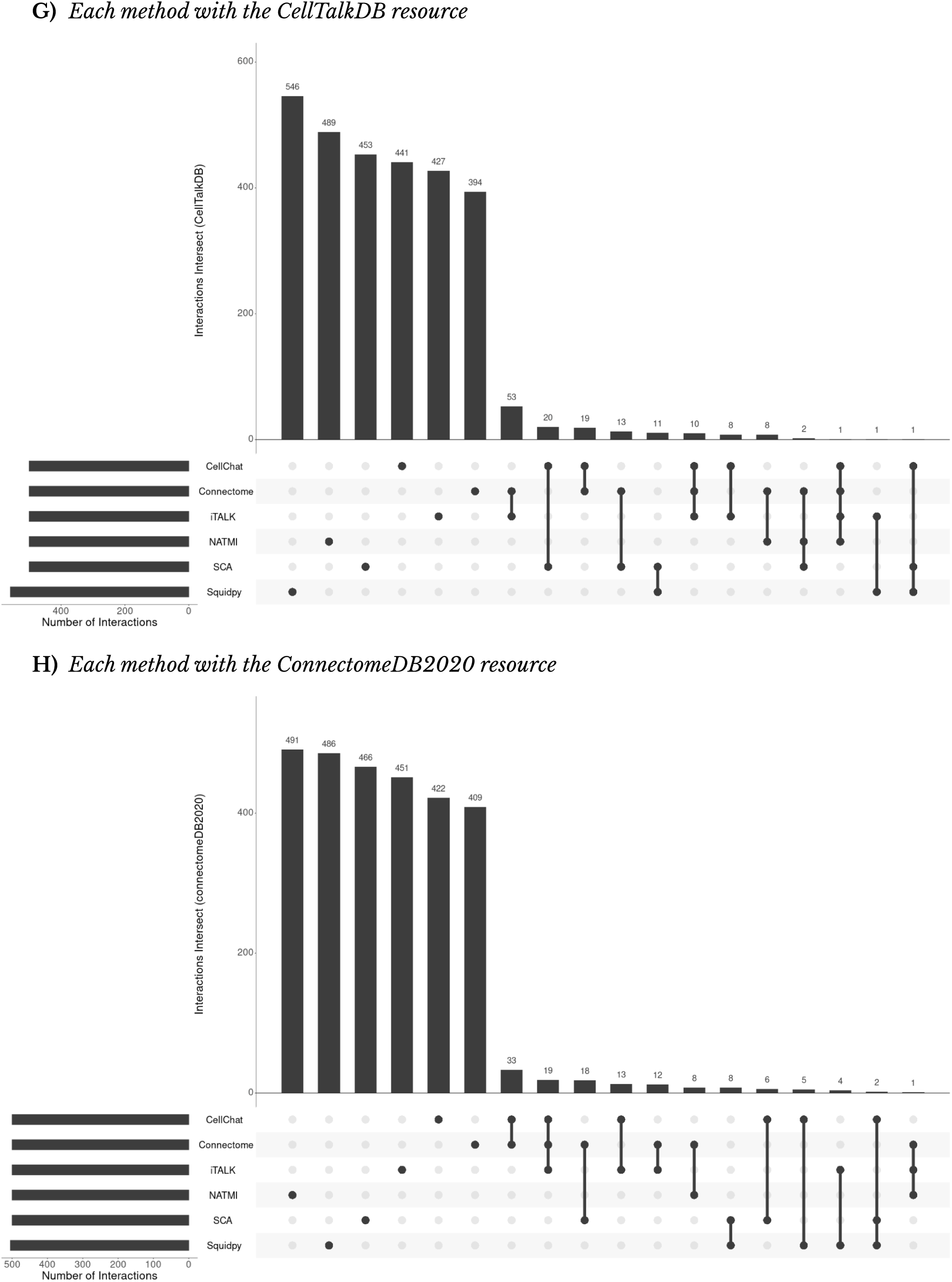
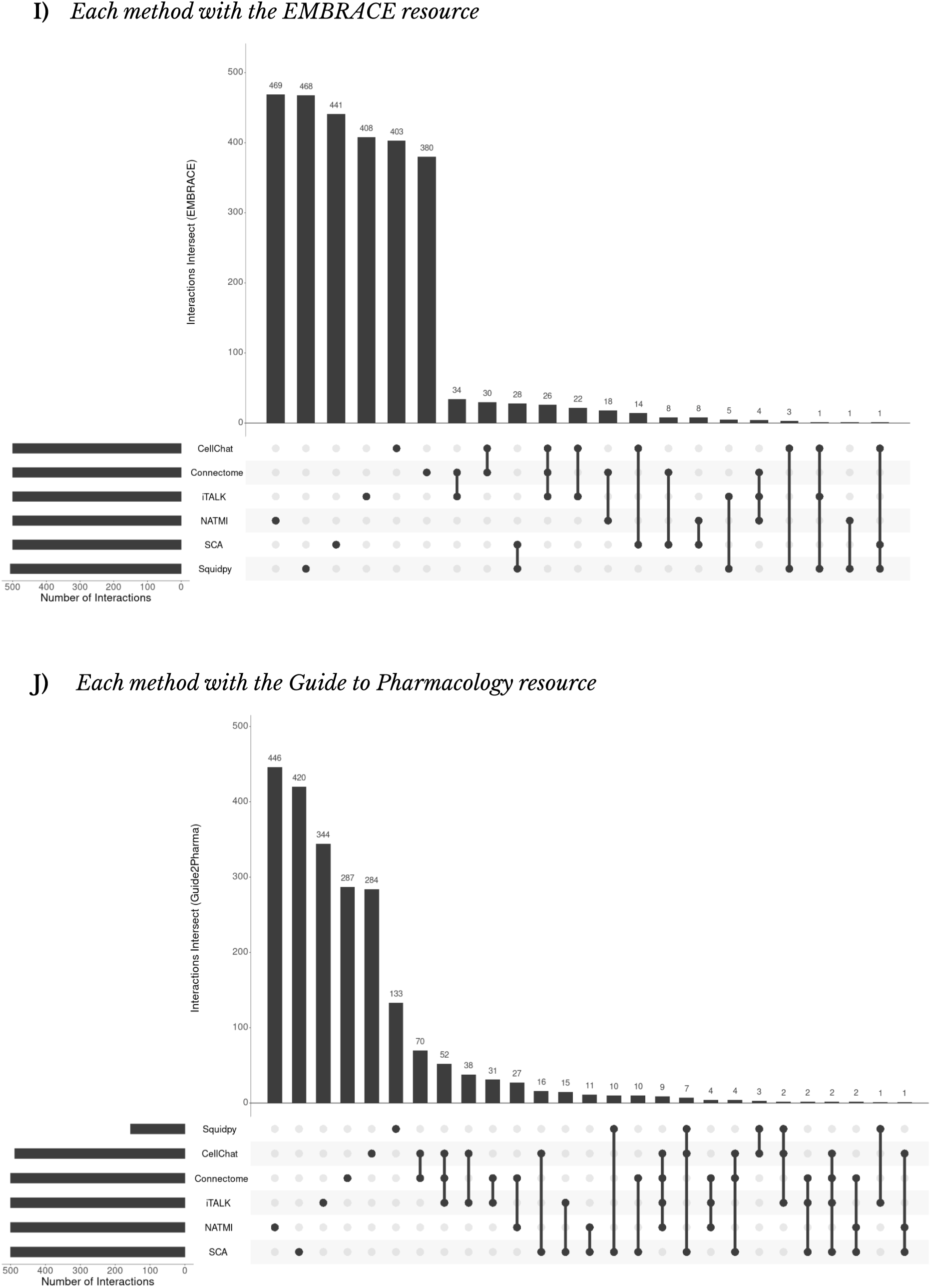
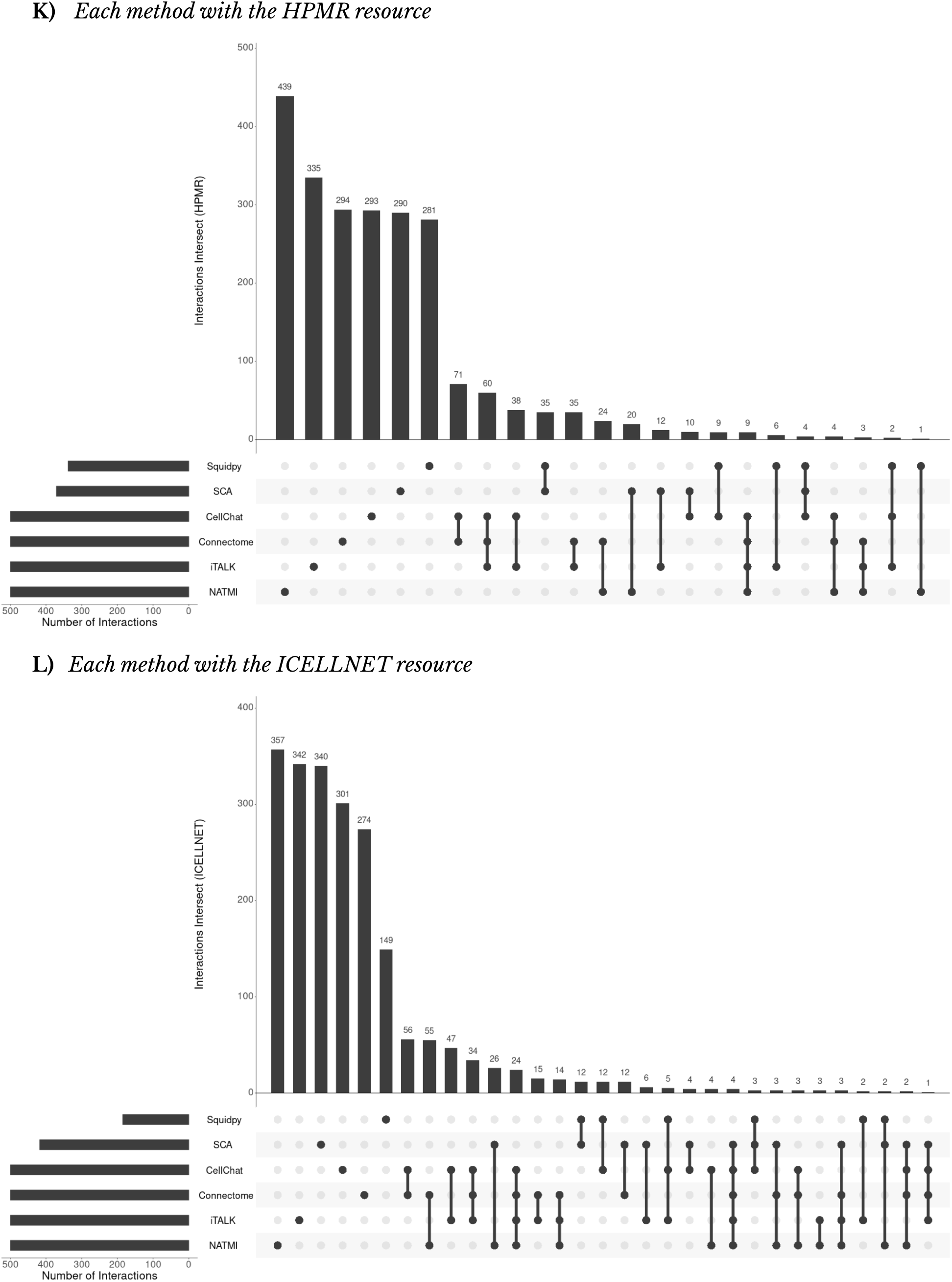
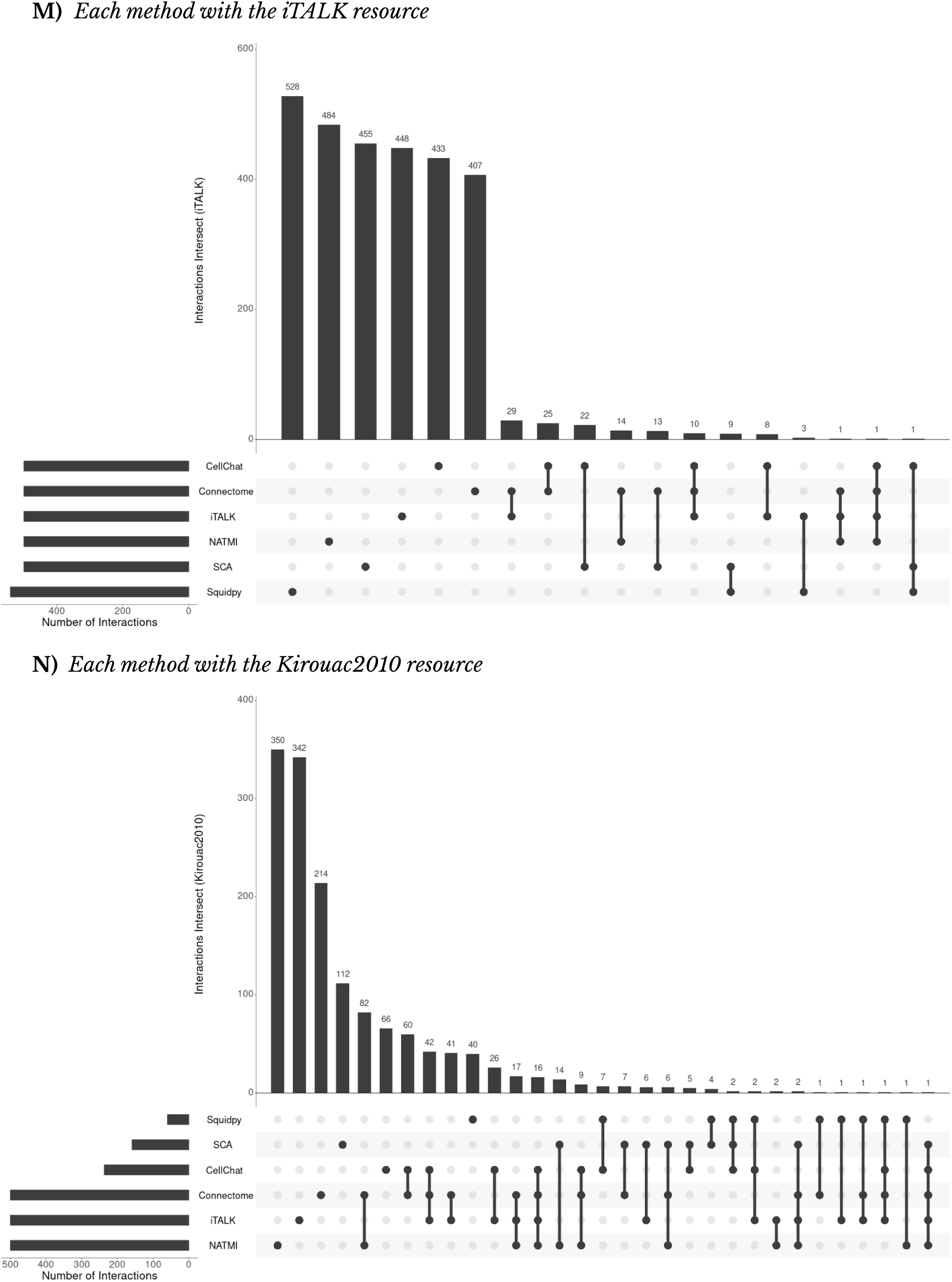
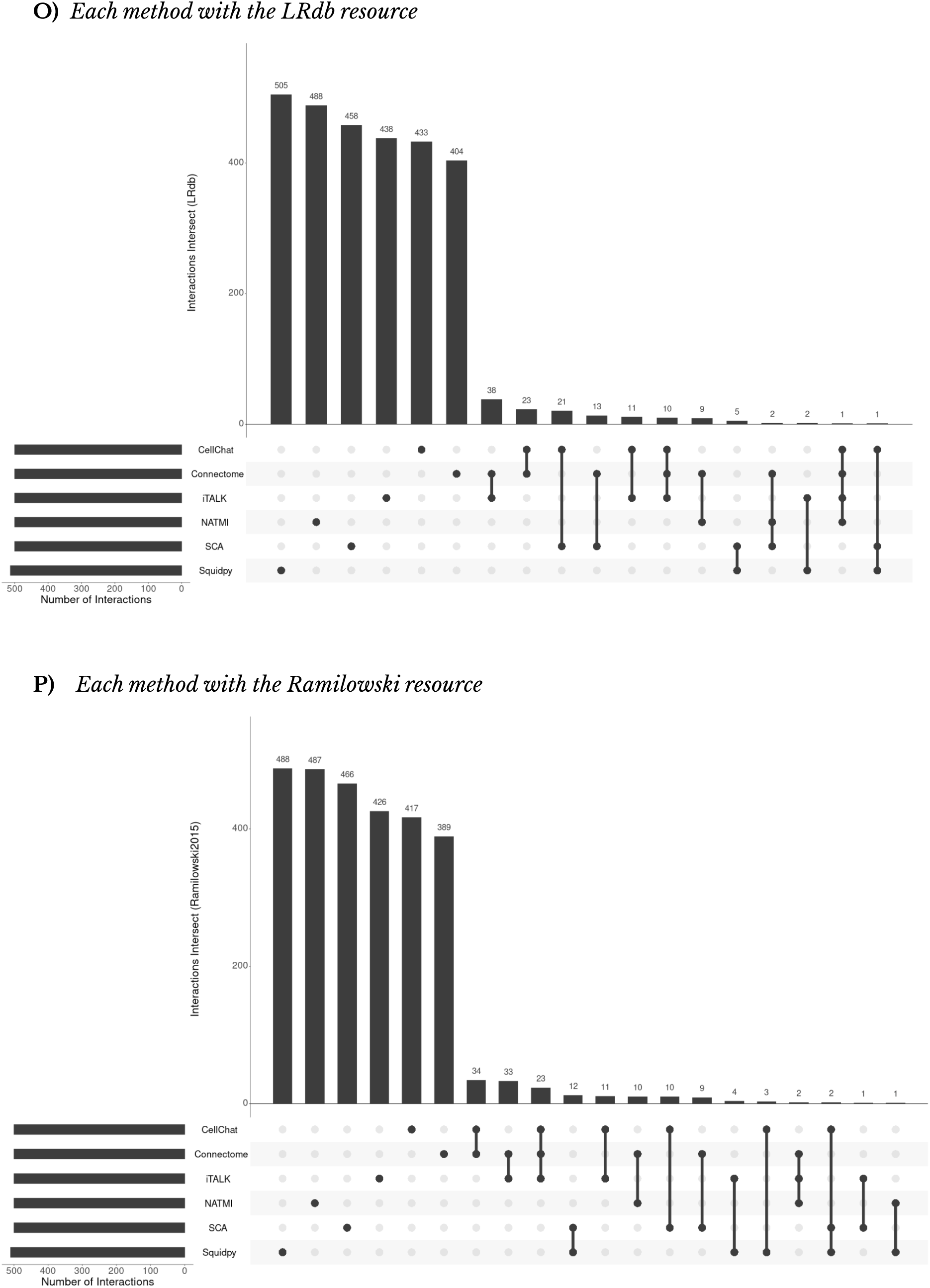
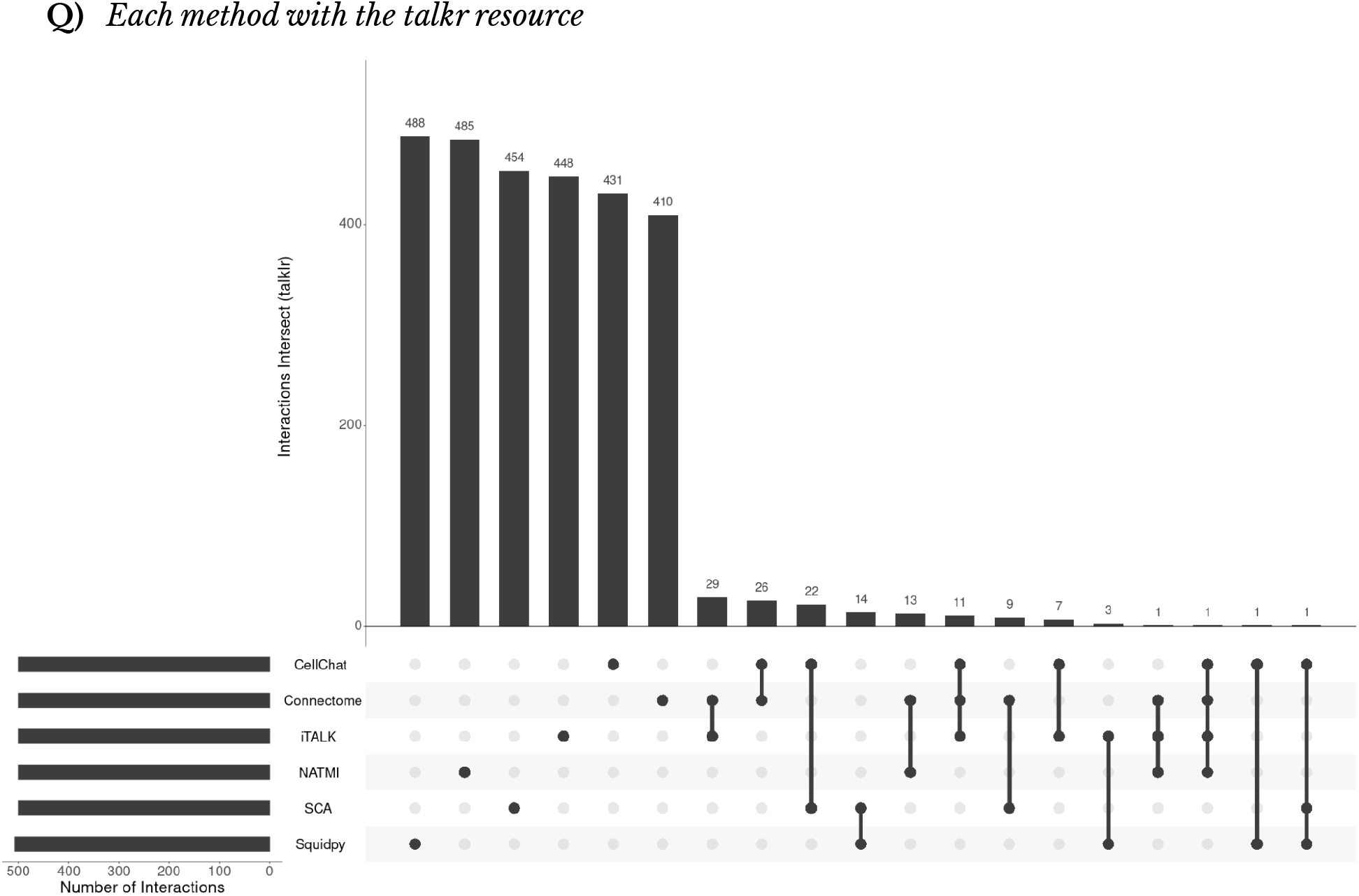
Upset plot showing overlap of most relevant interactions for each method with the same resource

**Supplementary Figure S15.**
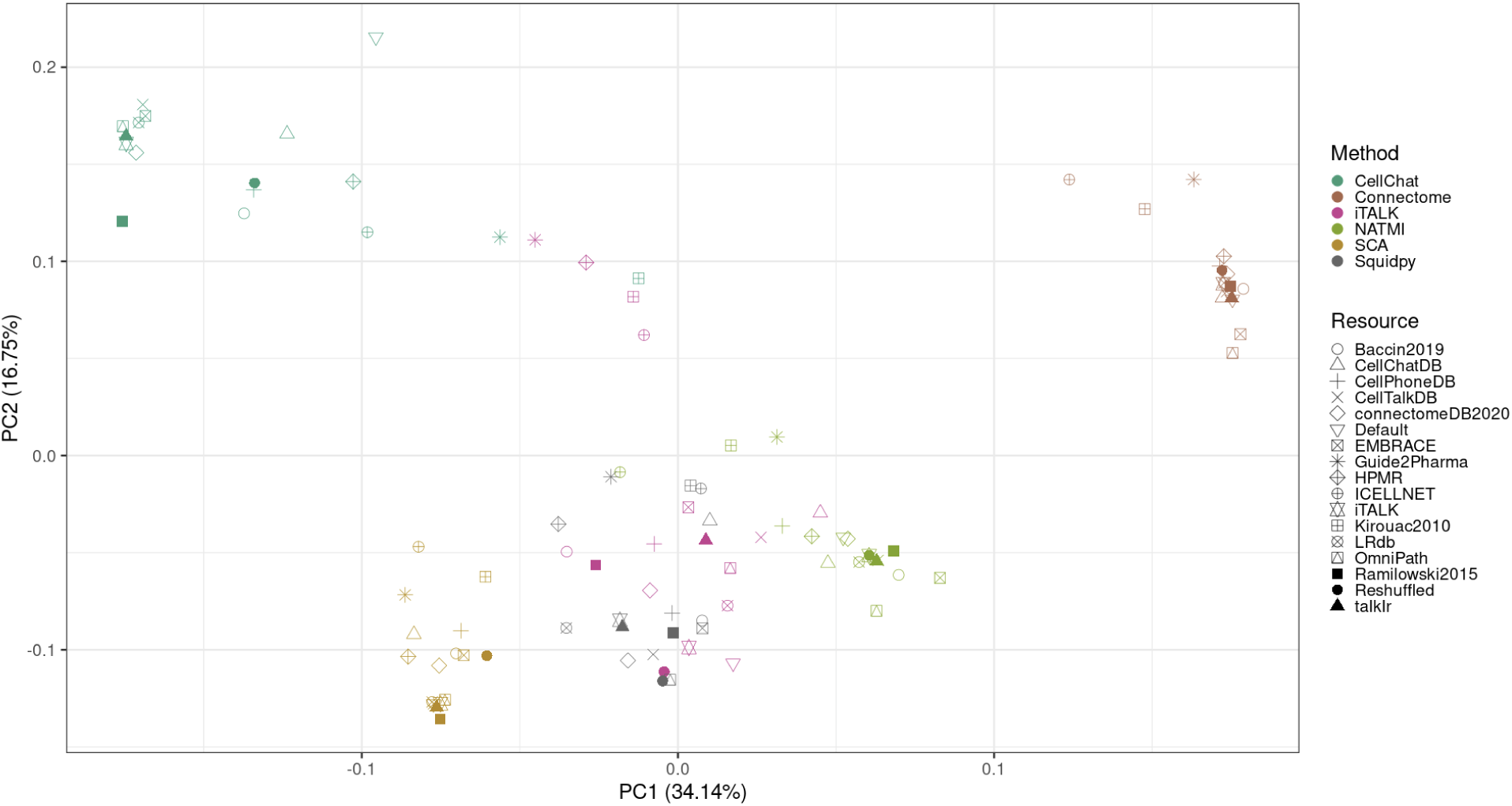
PCA of normalized average interaction rank frequencies per cell pair

**Supplementary Figure S16.**
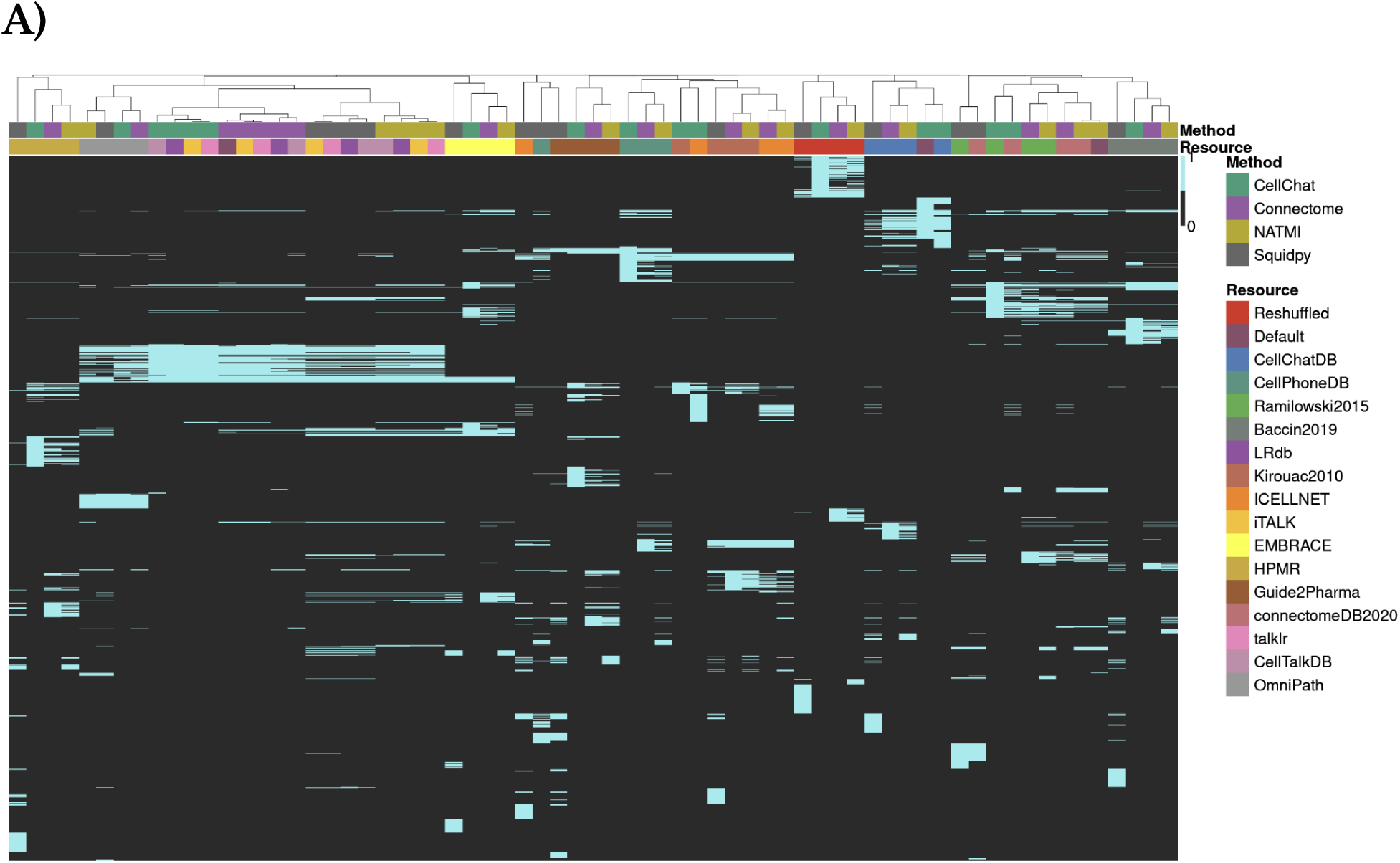
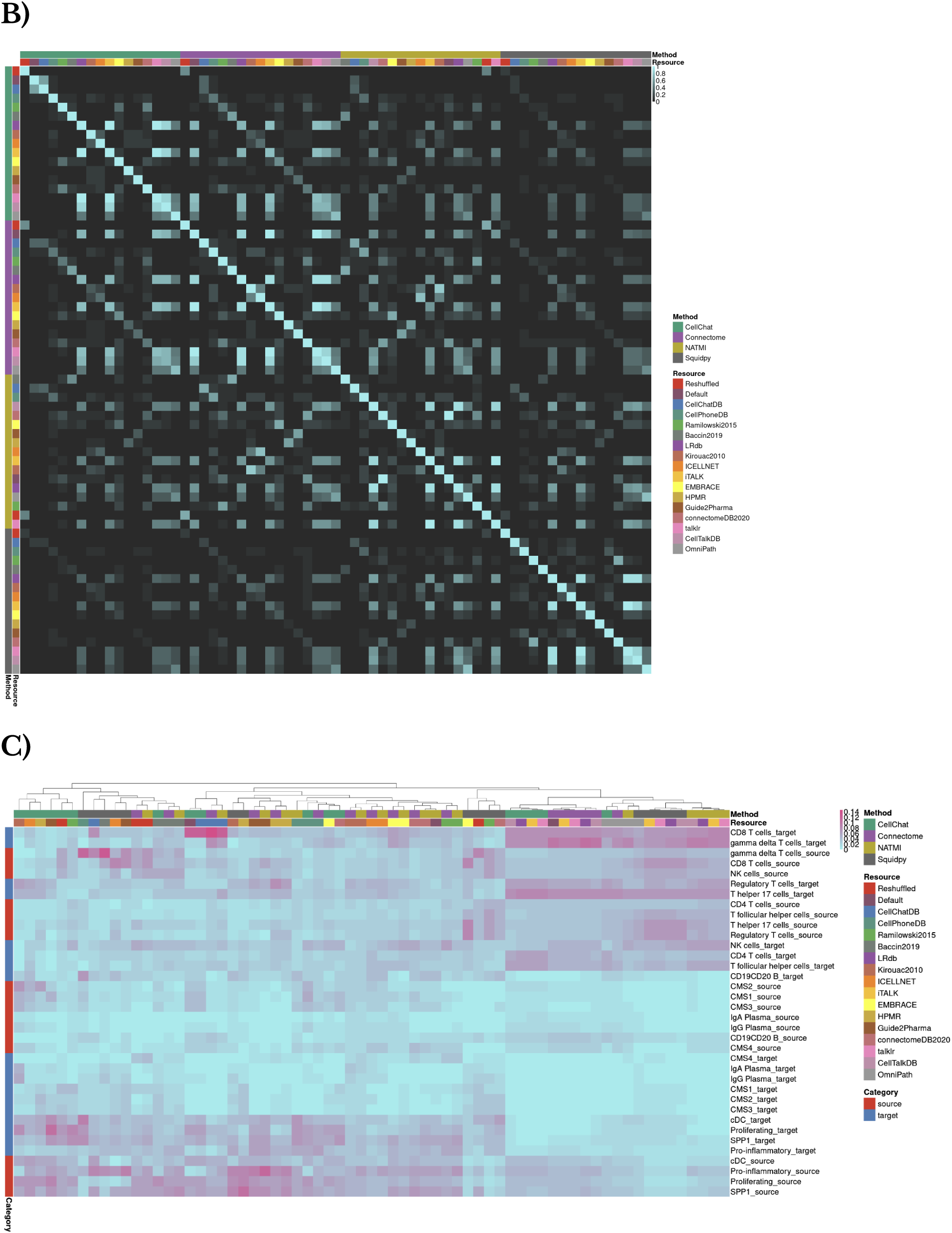
Cluster-unspecific communication agreement. **A)** Overlap in 500 highest ranked interactions between different combinations of methods and resources. **B)** Similarity among the highest ranked interactions for each method-resource combination, as measured by Jaccard index. **C)** Activity per Cell type, inferred as the proportion of interaction edges that stem from Source Cell types or lead to Target Cell types in the highest ranked interactions.

**Supplementary table 1.** Unique and shared Transmitters, Receivers, and interactions in each resource. We defined unique and shared interactions, receivers and transmitters between the CCC resources if they could be found in only one or at least two of the resources, respectively.

| Resource | Transmitters | Receivers | Interactions |
| --- | --- | --- | --- |
| Baccin2019 | 10.29% | 7.98% | 10.52% |
| CellChatDB | 11.73% | 17.17% | 50.29% |
| CellPhoneDB | 5.55% | 14.31% | 15.27% |
| CellTalkDB | 3.70% | 6.93% | 6.28% |
| ConnDB2020 | 7.41% | 6.04% | 9.14% |
| EMBRACE | 1.16% | 0.00% | 3.42% |
| GuidePharm | 0.69% | 0.41% | 5.89% |
| HPMR | 10.56% | 4.67% | 15.84% |
| ICELLNET | 1.38% | 2.53% | 3.64% |
| iTALK | 0.00% | 0.00% | 0.08% |
| Kirouac2010 | 2.84% | 0.00% | 9.33% |
| LRdb | 0.88% | 0.00% | 1.61% |
| OmniPath | 16.28% | 4.29% | 45.70% |
| Ramilowski | 0.00% | 0.00% | 0.00% |
| talklr | 0.00% | 0.00% | 0.00% |
| Total | 5.30% | 4.57% | 16.77% |

**Supplementary table 2.** Description of existing resources for measuring cell–cell communication. Formatted Korean CRC data set cell type counts and full names.

| Resource | Further curation | Sources | Interactions* |
| --- | --- | --- | --- |
| Baccin2019(a)<sup>22</sup> | Murine identifiers (only),<br>Multimeric complexes | Ramilowski2015, KEGG<br>reactome database, Literature | 1978 (1418) |
| CellChatDB<sup>13</sup> | Multimeric complexes, 229<br>signaling pathway families,<br>agonists and antagonists,<br>co-receptors, localisations,<br>Murine identifiers | KEGG, Literature | 2551 (2551) |
| CellPhoneDB<sup>7,25,26</sup> | Multimeric complexes,<br>intercellular communication<br>roles | Guide2Pharma, I2D, IMEx,<br>InnateDB, IntAct, MatrixDB,<br>MINT, UniProt, Literature | 1397 (1312) |
| CellTalkDB<sup>20</sup> | Murine identifiers | STRING, Literature | 3398 (3390) |
| ConnectomeDB2020<sup>10</sup> | - | Ramilowski2015,<br>CellphoneDB, Baccin2019,<br>LRdb, ICELLNET, Literature | 2293 (2264) |
| EMBRACE(a)<sup>61</sup> | Murine identifiers | Ramilowski2015 | 1710 (1489) |
| Guide2Pharma(b)<sup>31</sup> | - | Literature | 740 (662) |
| HPMR<sup>37</sup> | - | PubMed, GenBank | 527 (461) |
| ICELLNET<sup>21</sup> | Multimeric complexes,<br>Signalling families,<br>Cytokine-focus | STRING, Ingenuity, BioGRID,<br>Reactome, CellPhoneDB | 380 (371) |
| iTALK (c) <sup>5</sup> | Ligand categories | Ramilowski2015, HPMR,<br>IUPHAR-DB, Graeber2001,<br>Griffith2014, Cameron2003,<br>Zhou2017, Auslander2018 | 2648 (2565) |
| Kirouac2010<sup>38</sup> | - | Literature, COPE | 270 (150) |
| LRdb<sup>11</sup> | - | cellsignal.com,<br>Ramilowski2015,<br>Guide2Pharma, HPMR,<br>HPRD, Reactome, UniProt,<br>Literature | 3251 (3226) |
| Ramilowski2015<sup>32</sup> | - | DLRP, HPMR, IUPHAR,<br>HPRD, STRING, Literature | 1894 (1888) |
| talklr<sup>36</sup> | - | - | 2422 (2411) |
| OmniPath#<sup>2</sup> | Combines data from more than<br>100 resources and contains<br>protein-protein and gene<br>regulatory interactions,<br>enzyme-PTM relationships,<br>multimeric complexes, protein annotations, and a<br>CCC-dedicated composite DB | Composite resource<br>combining all of the CCC<br>dedicated resources listed<br>here, along with some<br>additional interactions. | 6103 |
| # All the resources above were retrieved from the OmniPath database (https://omnipathdb.org/). We also refer to the composite CCC resource presented here as OmniPath. The OmniPath presented in the analyses we filtered according to: **i)** we only retained interactions with literature references, **ii)** we kept interactions only where the receiver protein was plasma membrane transmembrane or peripheral according to at least 30% of the localisation annotations, and **iii)** we only considered interactions between single proteins<br>* The number of original interactions for each CCC dedicated resource covered in OmniPath is shown in brackets.<br>(a) Translated from murine identifiers to human, which accounts for the lower number of obtained interactions.<br>(b) Kept only the unique human-annotated interactions between transmitter and receiver proteins.<br>(c) Duplicates present in the original resource were excluded when imported via OmniPath |  |  |  |

**Supplementary table 3.** Formatted Korean CRC data set cell type counts and full names.

| Cell type | Cell subtype | Complete Name | Cell Count |
| --- | --- | --- | --- |
| B cells | CD19CD20 B | B cells_CD19+CD20+ | 2,049 |
| T cells | CD4 T cells | CD4+ T cells | 3,980 |
| T cells | CD8 T cells | CD8+ T cells | 4,647 |
| Myeloids | cDC | Conventional Dendritic Cells | 353 |
| Epithelial cells | CMS1 | Consensus Molecular Subtype 1 | 1,201 |
| Epithelial cells | CMS2 | Consensus Molecular Subtype 2 | 10,771 |
| Epithelial cells | CMS3 | Consensus Molecular Subtype 3 | 5,486 |
| Epithelial cells | CMS4 | Consensus Molecular Subtype 4 | 11 |
| T cells | gamma delta T cells | $\gamma\delta$ T cells | 219 |
| B cells | IgA Plasma | IgA+ Plasma Cells | 180 |
| B cells | IgG Plasma | IgG+ Plasma | 1,661 |
| T cells | NK cells | Natural Killer Cells | 948 |
| Myeloids | Pro-inflammatory | Pro-inflammatory Macrophages | 2,325 |
| Myeloids | Proliferating | Proliferating Macrophages | 165 |
| T cells | Regulatory T cells | Regulatory T cells | 2,943 |
| Myeloids | SPP1 | SPP1+ Macrophages | 3,096 |
| T cells | T follicular helper cells | T follicular helper cells | 548 |
| T cells | T helper 17 cells | T helper 17 cells | 1,961 |

**Supplementary Note 1**. *Protein localisation to categorize CCC*

To estimate the localisation distributions of transmitters and receivers as well as to categorize CCC interactions according to signalling categories we obtained protein subcellular localisation annotations via OmniPath^2^. These annotations were gathered from sources such as UniProt, the Cell Surface Protein Atlas^62^, and Membranome^63^. The localisation annotations were then filtered according to a consensus threshold (**4.1. Descriptive analysis of resources**). We then used the localisations of transmitters and receivers to approximate the categories of interactions. The largest part of interactions were those between secreted proteins targeting transmembrane proteins (S -> T), which we referred to as the secreted signalling category in Figure 3. Further, we attributed interactions between and within the transmembrane and peripheral plasma membrane proteins (T -> T, P -> T, T -> P, P -> P) to intercellular signalling events that require physical contact between cells.

As a consequence of the protein localisation annotation process, some annotations were expected to be unsuitable in the context of CCC signalling. For example, a ligand can be annotated as both secreted and membrane-bound, depending on the context of the observation. This was the case for EFNA1, a membrane-bound ligand which binds to the EPH receptors, also observed to be released as a soluble protein in breast adenocarcinoma cells^64^. Moreover, splicing variants of the same protein can have different subcellular localisations, with one variant being membrane-bound and the other secreted. For instance, FGF17 binds to membrane-bound FGFR2, but since FGFR2 also has secreted isoforms (UniProtKB - P21802), this interaction can be mislabeled as an interaction between two secreted proteins. These misannotations in the context of CCC interactions made up only a small proportion of all annotations and were grouped into the “Other” category (Figure 3), which represented interactions which did not fit as secreted or direct-contact signalling (T->S; S->S; P->S).

**Supplementary Note 2.** *Protein Complexes*

We further assessed the predicted proportions of interactions containing complexes from CellChat and Squidpy using Baccin, CellChatDB, CellPhoneDB, and ICELLNET resources. This analysis showed that the proportion of complexes among the highest ranked 500 hits for CellChat ranged from 1.8% (with ICELLNET) to 23.0% (with the original CellChatDB) and that of Squidpy ranged from 9.7% with ICELLNET) to 38.1% (with CellChatDB).

**Supplementary Note 3**. *Cluster Specificity and Method Dissimilarity*

As a consequence of the disagreement between methods in regards to the most actively communicating cell types, we reasoned that a possible cause was the different approaches used to assign cell cluster specificity to the interactions. To this end, we conducted the same analyses presented in the main text, but instead using the measures from each method which do not explicitly reflect the cell-type specific communication (i.e. Squidpy means; unfiltered CellChat probabilities; Connectome.weight.norm; NATMI.edge.avg.expr) (Table 1). Since SingleCellSignalR and iTALK provide a single scoring system, they were excluded from this analysis. We observed an increase in the agreement between methods (Supp. Figure 16A), as the mean Jaccard index when using the same resource with different methods ranged from 0.277 to 0.618 (mean = 0.404) (Supp. Figure 16B). The overlap between these methods when using the same resource was hence considerably higher than that observed when using cluster-specific measures (mean = 0.0247). On the contrary, the mean Jaccard index per method when using the same resource remained relatively unchanged when compared to the scoring systems that reflect cell cluster specific communication (0.118 for non-specific measures versus 0.167 for specific). Moreover, analogously to the agreement analysis, we used the cluster-unspecific measures to estimate the active cell types. As a result, methods were observed to largely agree in terms of the most active cell types (Supp. Figure 16C). Thus, this analysis suggests that the distinct approaches used to assign cell cluster specificity to the interactions explain some of the disagreement between methods for our dataset. Furthermore, when using the cluster-unspecific measures, the differences in resources were the main source of dissimilarity between the results.

**Supplementary Note 4**. *Single-cell CCC Benchmark Directions*

As a consequence of the observed disagreement in the results obtained when using different methods and resources, we argue that an appropriate benchmark is paramount for the future development of the CCC inference field. Some effort has already been directed to the assessment of different methods and specific directions were already proposed^19^. To this end, we also share our current benchmark ideas.

### I. Associations between CCC activity and Spatial-Adjacency

#### Assumptions

Cell clusters that are spatially adjacent should be communicating more actively than those that are spatially distant; Confining CCC inference to spatial adjacency should reduce false positives.

#### Limitations

Difficult to distinguish cell-cell communication and cellular program coregulation.

#### Examples

This approach was already used as a way to validate some methods^13, 18^, while other methods explicitly take spatial information into account for CCC inference^19^. Another example is confining CCC inference to cells that are expected to be in close contact, e.g. according to co-localizing cells in visium spots to reduce false positive interactions^25^. In a similar way, 10x Visium data can be used to identify cell types that are known to be co-located in visium spots, and are hence in close contact.

#### Metrics

A benchmark focused on the relationship between Cell Pair Activity*^1^ and Cell Distance*^2^ composed by three main steps:

(1) Cell Pair Activity reported by different methods (**\*1**);
(2) Cell-cell Spatial Distance or Colocalization (**\*2**);
(3) Relationship between CCC method output and distance (**\*3**).

**\*1**. Number of Inferred interactions between Cell Clusters; Average Cell Inference Ranks per Communicating Cell Types

**\*2**. Physical distance, measured by Euclidean distance between the closest cell types, was already reported to be an appropriate proxy of cell pair communication activity^65^. Other measures can be the neighbourhood enrichment or spatial co-occurrence of cells^24, 66^. An alternative approach would be to discretise distance according to e.g. spatially-adjacent and spatially-distant cell types^13, 18^.

**\*3**. Correlation or Regression Coefficients, or any other measure used as a proxy of the relationship between the two variables.

### II. Data-driven Inference of Spatial Covariance to explain Transmitter-Receiver interactions

#### Assumptions

Receiver and transmitter gene expression covariance with spatial distance is a proxy of CCC events.

#### Limitations

Difficult to distinguish cell-cell communication and cellular program coregulation; Possibly biased towards CCC events in which the transmitter and receiver regulate each other’s expression.

#### Approach

**1)** The expression of a receiver is spatially explainable by the expression of a transmitter and vice versa. Thus, a threshold signifying conserved spatial gene regulation between transmitters and receivers (with e.g. mistyR^67^) can be used to define putative true positive interactions.
**2)** Downstream signalling models to explain transmitter and receiver activity.

Some tools already utilize downstream signalling as an attempt to better model CCC interactions^15, 16, 18^. In a similar way prior knowledge of downstream transmitter and receiver activity models^15^, can be used to spatially explain the activity of transmitter and receiver. Alternatively, one can build naïve protein-protein interaction models from existing databases^2^.

#### Metrics

Methods’ (and Resources’) coverage of the spatially explainable CCC events. In other words, we expect a method to assign preferentially high ranks to spatially explainable transmitter-receiver interactions.

AUROC can be calculated according to different thresholds of spatial covariance for transmitter/receiver genes involved in CCC interactions. Thus, a reliable tool and resource should be able to pick up spatial covariance better than a randomized resource, and better than a resource composed of genes that are not explainable by space (e.g. housekeeping genes).

